# Spatiotemporal transcriptomic profiling reveals the dynamic immunological landscape of alveolar echinococcosis

**DOI:** 10.1101/2024.04.04.577902

**Authors:** Zhihua Ou, Li Li, Peidi Ren, Ting-Ting Zhou, Fan He, Jialing Chen, Huimin Cai, Xiumin Han, Yao-Dong Wu, Jiandong Li, Xiu-Rong Li, Qiming Tan, Wenhui Li, Qi Chen, Nian-Zhang Zhang, Xiuju He, Wei-Gang Chen, Yanping Zhao, Jiwen Sun, Qian Zhang, Yan-Tao Wu, Yingan Liang, Jie You, Guohai Hu, Xue-Qi Tian, Sha Liao, Bao-Quan Fu, Ao Chen, Xue-Peng Cai, Huanming Yang, Jian Wang, Xin Jin, Xun Xu, Wan-Zhong Jia, Junhua Li, Hong-Bin Yan

## Abstract

Alveolar echinococcosis (AE) is caused by the invasive growth of the metacestodes of *Echinococcus multilocularis* (*E. multilocularis*). The early diagnosis, management, and treatment of AE remains challenging. Herein, we integrated bulk RNA-seq, scRNA-seq, and ST technologies to reveal the immune characteristics both spatially and chronologically in *E. multilocularis* infected mouse liver. An unprecedented high-resolution spatial atlas of the *E. multilocularis* infection foci was obtained, revealing the pivotal role of neutrophils, *Spp1*^+^ monocyte-derived macrophages (MoMFs), and fibroblasts in AE progression. We observed continuous recruitment of neutrophils and macrophages into the infected tissues, indicating prolonged immune stimulation by the growing metacestodes. However, their spatial distribution patterns and functions changed during the infection cause. At the early infection stage, *Il1b*^hi^ neutrophils aggregated in the lesion displayed the highest pattern-recognition receptor activity and may kill the metacestodes through NETosis, but their death in the infection foci might also trigger immune suppression. Meanwhile, *Spp1*^+^ MoMFs attached to the protoscoleces to clear the infection. At the late infection stage, the neutrophils failed to infiltrate into the lesion although their continuous recruitment into the infected liver. Even though the *Spp1*^+^ MoMFs enclosing the microcysts of *E. multilocularis* may retain their pathogen-killing ability, they may also promote fibrosis and angiogenesis through interaction with fibroblasts. The pro- and anti-inflammatory phenotype balance of these immune cells may be associated with the transition of parasite control strategy from “active killing” to “negative segregation” by the host. Our findings identify key molecular traits involved in AE development, which may benefit the treatment of echinococcosis.

## INTRODUCTION

Alveolar echinococcosis (AE) is a zoonotic parasitic disease caused by *Echinococcus multilocularis* (*E. multilocularis*), one of the 20 targeted neglected diseases for control or elimination in impoverished areas by The World Health Organization (WHO) roadmap 2021-2030 ^1^. It is mainly prevalent in the Northern Hemisphere and has a high incidence in the pastoral areas of Asia, North America and Europe ^2,3^. Human infection is caused by the ingestion of water and food contaminated by the eggs of *E. multilocularis,* which primarily occurs in the liver. The infiltrative and oppressive tumor-like growth of metacestodes is the main cause of the disease, which can damage blood vessels, bile ducts, and adjacent organs. The host organs are concurrently affected by the mechanical compression, toxin stimulation, and invasive growth of the metacestodes ^4–6^. The untreated patients may reach a mortality rate higher than 90% ^7^. Imaging diagnostic techniques for AE included computed tomography (CT) ^8,9^, magnetic resonance imaging (MRI) ^10,11^ and ultrasound ^12,13^, but the imaging features of AE patients in the early stage of infection are lack of specificity. Thus, most of the diagnosed AE patients have entered the later stage of the disease and missed the best opportunity for surgical operation. AE lesions are usually found in the right liver, invading bile ducts and hepatic veins in advanced cases, making liver resection impossible due to the risk of uncontrolled bleeding and the high demands on surgical techniques ^14^. Radical surgery can only benefit a small number of patients with AE on the left lobe. The available anti-echinococcosis drugs, such as albendazole and mebendazole, are only parasitostatic, requires prolonged administration and sometimes have serious side effects ^15^. Therefore, there is an urgent need to enhance the understanding of the cellular and molecular interactions involved in AE progression in order to bring new insights into echinococcosis prevention and treatment ^16^.

Liver pathogenesis caused by *E. multilocularis* infection is manifested by helminthic microcysts surrounded by immune cells and fibrotic tissues ^14^. After infection, lots of the immune cells, especially neutrophilic cells, are present in the infection foci, secreting inflammatory cytokines to fight against the parasites ^17^. However, the exact cell composition in the infection tissues and the cellular structural dynamics during sustained infection remain obscure. In the late infection stage, the upregulation of exhaustion-related genes such as *NFKBIA*, *PD-1*, *CTLA4*, *TIGIT,* and *TIM-3* transforms the immune cells including *CD4^+^* and *CD8^+^* T cells into an immune inhibition phenotype, facilitating the persistent propagation of the metacestodes in the host ^18–20^. Neutrophils are the first leukocytes recruited to eliminate invading pathogens through multiple mechanisms such as NETosis and phagocytosis ^21^. However, the role of neutrophils in the long-term development of AE remains poorly understood.

Based on histopathological observations and the detection of known immune-related genes and proteins, a general map of the liver injury caused by *E. multilocularis* infection can be obtained. However, a high-resolution spatial landscape of the infection foci and the cell-cell interaction mechanism driving disease progression remain to be elucidated. To solve this problem, single-cell RNA sequencing (scRNA-seq) can be applied to acquire the transcriptomes of single cells after tissue dissociation, which can reveal the changes in cell types and cell states during parasite infection ^18,19^. Meanwhile, the spatial transcriptomics (ST) technology can reveal the cell-cell interactions in specific pathological niches by capturing tissue transcriptomes *in situ*. Herein, we integrated data generated by multiple transcriptomic technologies, including bulk RNA-seq, scRNA-seq, and ST technologies, to reveal the cellular compositions and molecular interaction characteristics both spatially and chronologically in *E. multilocularis* infected liver based on a murine model. Our results unveiled an unprecedented high-resolution spatial atlas of the *E. multilocularis* infection foci and the functional roles of neutrophils, *Spp1^+^* macrophages, and fibroblasts during disease progression, enhancing our understanding of the pathogenesis of AE.

## RESULTS

### The cellular composition of the AE lesions in mouse liver

Clinical observations based on H&E and IHC staining have revealed the aggregation of multiple types of immune cells in the infection foci of *E. multilocularis* in the liver. Yet the detailed spatial organization of the immune cells around the infection foci and the communication mechanisms between the neighboring cells remain to be elucidated. Herein, we used a high dose of *E. multilocularis*, 2,000 protoscoleces diluted in PBS, to infect BALB/c mice. Liver tissues were collected at multiple timepoints to track the pathological progress after infection (Figure 1A). Using Stereo-seq technology ^22^, we obtained the spatial gene expression profiles of 14 mouse liver samples, including 1 from the control mice, 5 from 4 days post inoculation (dpi), 3 from 8 dpi, 3 from 15 dpi, 1 from 37 dpi, and 1 from 79 dpi (Figure S1A; Table S1). The Stereo-seq chips were 1 x 1 cm^2^ in size, and the capture spots had a diameter of 220 nm and a center-to-center distance of 500 nm. We used a spatial unit of bin100 for the overall analysis of the 14 Stereo-seq chips to ensure sufficient gene types in the bins of each chip. A bin100 represents a square area with a side length of 100 spots, covering a region of 49.72 x 49.72 μm ^22^. The number of gene types per bin100 ranged from 349 to 2,250, with a median number of 1,300 (Table S1). The dominant cell types of each bin were predicted using the liver cell atlas contributed by Guilliams et al as reference ^23^, which contained most of the cell types in the mouse liver, and the annotation results were further supported by the expression of marker genes (Figure S1B). We identified eight major cell types in the tissues from infected and healthy mouse livers (Figures 1B, 1C, S1A, and S1B), including hepatocytes (*Alb*, *Apoc1*, *Glul*, and *Cyp2e1*), fibroblasts (*Col1a1*, *Col1a2*, and *Timp1*), neutrophils (*S100a8*, *S100a9*, *Retnlg*, *Nfkbia*, *Il1b*, and *G0s2*), *Spp1*^+^ monocyte-derived macrophages (*Spp1*^+^ MoMFs, expressing *Cd68*, *Cd14*, *Spp1*, *Csf1r*, *Adgre1*, and *Itgam*), monocyte-derived Kupffer cells (MoKCs, expressing *Clec4f*, *Cd5l*, and *Vsig4*), cholangiocytes/*Spp1*^+^ cells (*Spp1*), plasma cells (*Igkc*, *Jchain*, *Ighm*, and *Mzb1*), and hepatic stem/progenitor cells (HsPCs, expressing *Epcam*, *Krt19*, and *Sox9*). Among them, neutrophil was the major cell type in the granulomatous structure of the AE lesion (Figures 1B and S1A). Temporal comparison of the Stereo-seq data showed that the populations of four cell types, i.e., neutrophils, *Spp1*^+^ MoMFs, plasma cells, and fibroblasts were abundantly expanded after infection (Figure 1C). We further conducted a bulk RNA-seq experiment to identify the temporal dynamics of cell composition in mouse liver infected by *E. multilocularis*. Deconvolution of bulk RNA-seq data also revealed proportional changes in neutrophils, macrophages, plasma cells, and fibroblasts (Figures 1D and 1E), which were consistent with the trends observed in the spatial transcriptomic dataset. Based on the cell type annotation results, the spatial organization of AE lesions and associated cell-cell communication pathways present in disease progression can be further resolved.

**Figure 1.**
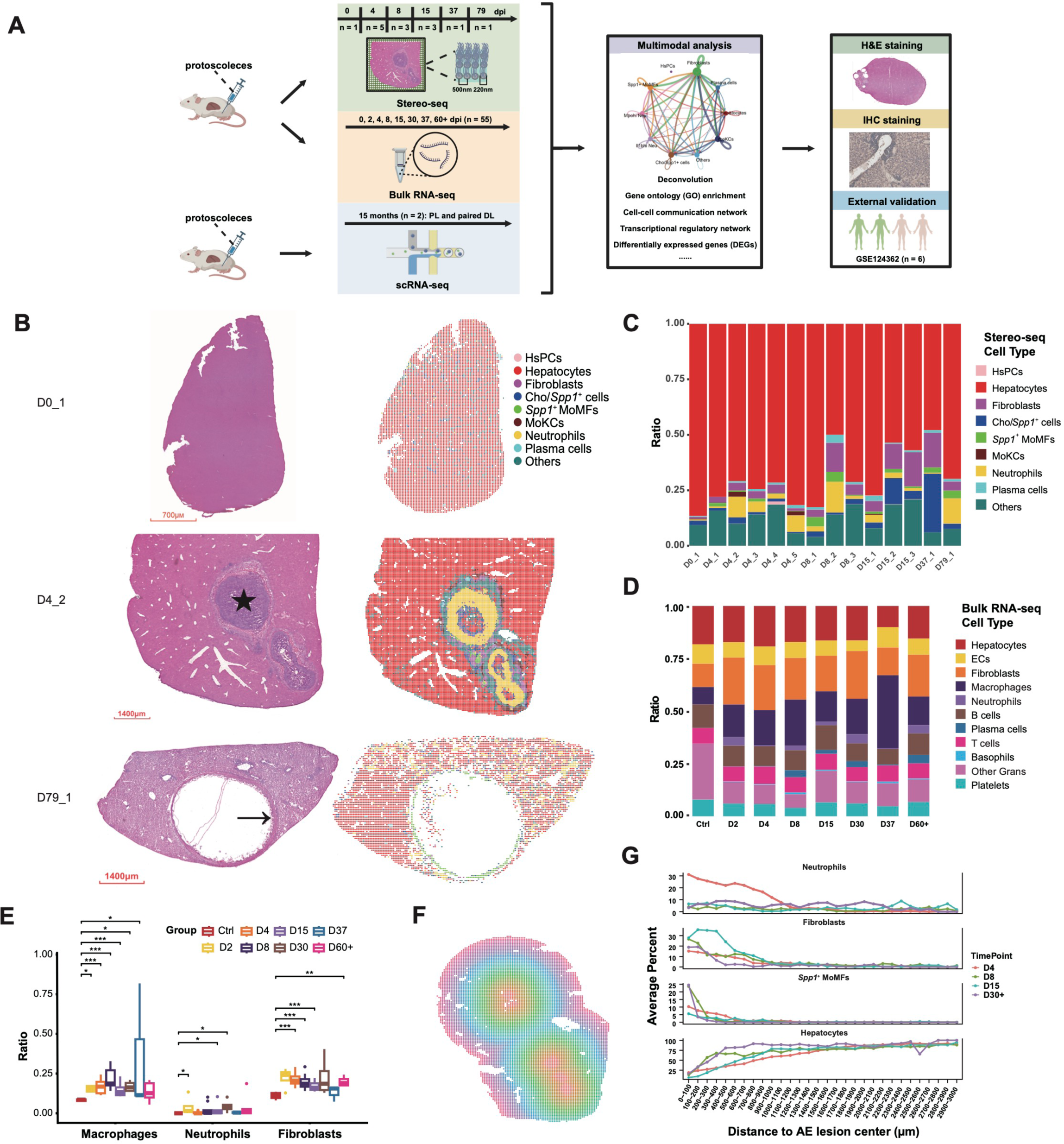
Spatiotemporal transcriptomic profiling of mouse liver infected with *Echinococcus multilocularis*. (A) Workflow of this study. n indicates the number of samples. (B) Cell type annotation results of three representative Stereo-seq slides. H&E staining was performed using the tissue section immediately adjacent to the tissue section used for Stereo-seq. The star mark indicates protoscoleces (PSCs), while the arrow indicates the microcyte of *Echinococcus multilocularis*. Analytical unit for Stereo-seq slides: bin100 (49.72 x 49.72 µm). D0_1: 0 day post infection, sample No.1; D4_2: 4 days post infection, sample No.2; D79_1: 79 days post infection, sample No.1. Cho: cholangiocyte; HsPC: hepatic stem/progenitor cell; MoMF: monocyte-derived macrophages. (C) Cell compositions of mouse liver samples collected at different timepoints based on Stereo-seq data. (D) Cell compositions of mouse liver samples collected at different timepoints (n = 55). The cell types were deconvoluted based on bulk RNA-seq data. EC: endothelial cell; Gran: granulocyte. (E) The number of macrophages, neutrophils, and fibroblasts was significantly increased in mouse liver after *E. multilocularis* infection. The cell types were deconvoluted based on bulk RNA-seq data (n = 55). (F) Schematic graph showing the layer distribution from the AE lesion center to distal region, based on Sample D4_2. (G) Gene Set Variation Analysis (GSVA) scores showing the distribution of neutrophils, fibroblasts, *Spp1^+^* MoMFs, and hepatocytes from the center to the distal region of the AE lesions of different timepoints.

### The spatial architecture of the AE lesions in mouse liver

While the Stereo-seq protocol was not optimal for capturing the mRNA of *E. multilocularis*, parasitic genes can still be detected in 13 Stereo-seq chips (Figure S1C). The worm genes were mainly expressed in the infection foci with protoscoleces (PSCs) or parasitic vesicles. To fully reveal the intricate spatial organization of the infection foci, we determined the lesion center based on parasitic gene expression or H&E staining observation. We further divided different spatial layers starting from the lesion center to the distal-lesion region based on a radius distance of 100 μm (Figures 1F and S2A) and calculated the proportions of cell types and GSVA (Gene Set Variation Analysis) scores of gene sets in each layer. The cell compositions exhibited significant differences as the distance changed (Figure 1G). The proportion of hepatocytes was the lowest within the lesion (Figures 1G and S2B), with few normal hepatic lobule structures in these regions in H&E staining images. As the distance from the parasitic center increased, the proportion of hepatocytes increased, along with an increase in GSVA scores of gene sets associated with normal liver functions, such as fatty acid metabolism and bile acid metabolism (Figure S3A). Results showed that neutrophils could swarm around the infection foci, covering a circle with a radius of 1,200 μm starting from the lesion center (Figures 1G and S2C), especially on 4 dpi, which was further confirmed by the high GSVA scores of functional pathways of neutrophils (Figure S3B). Meanwhile, fibroblasts were also dominant around the infection foci, which covered smaller space than neutrophils at the early stage but larger space at the later stage and overlapped the distribution area of neutrophils (Figures 1G, S1A, and S2D). The *Spp1*^+^ MoMFs were significantly enriched in the space closely adjacent to the infection foci (less than 600 μm) (Figures 1G and S2E), especially around the parasitic vesicles, as shown in a sample collected on 79 dpi (Figure 1B). These observations indicated pivotal roles of neutrophils, *Spp1*^+^ MoMFs, and fibroblasts during AE progression. The spatial atlas of the AE lesions provided physical clues for further investigation on the cell-cell interaction networks involved in the immune response and pathological damage induced by parasite infection.

### Cellular remodeling at the late stage of *Echinococcus multilocularis* infection

To elucidate the cellular composition of AE lesions at the late infection stage, we performed scRNA-seq using the distal-lesion (DL) and peri-lesion (PL) liver tissues from *E. multilocularis* infected mice (15 months post infection), obtaining a total of 39,199 single cells. Eleven cell types were annotated using a marker-based method (Table S2), including hepatocytes, fibroblasts, endothelial cells (ECs), neutrophils, other granulocytes (Other Grans), basophils, macrophages (MFs), B cells, plasma cells, T cells and platelets (Figures 2A and 2B). We further integrated the mouse liver atlas generated by Guilliams et al. with our scRNA-seq data to reveal the hepatic cellular changes between healthy and infected mice ^23^. Comparative analysis revealed significant increases in B cells, plasma cells, neutrophils, and fibroblasts in the infected tissues (DL and PL) compared to the healthy control group (Figure 2C). Additionally, the proportion of macrophages and fibroblasts significantly increased in the PL group compared to the DL group (Figure 2D). This revealed the dynamic cellular remodeling of the lesion microenvironment at a late infection stage of *E. multilocularis* in the mouse liver. We further used Tangram to integrate our scRNA-seq dataset with the Stereo-seq data, mapping gene signatures to the spatial dimension ^24^. The distribution of neutrophils and macrophages overlapped with the spatially annotated neutrophil-enriched and macrophage-enriched regions (Figure 2E), which mainly gathered around the AE lesion. Meanwhile, some neutrophils were scattered in other inflammation regions.

**Figure 2.**
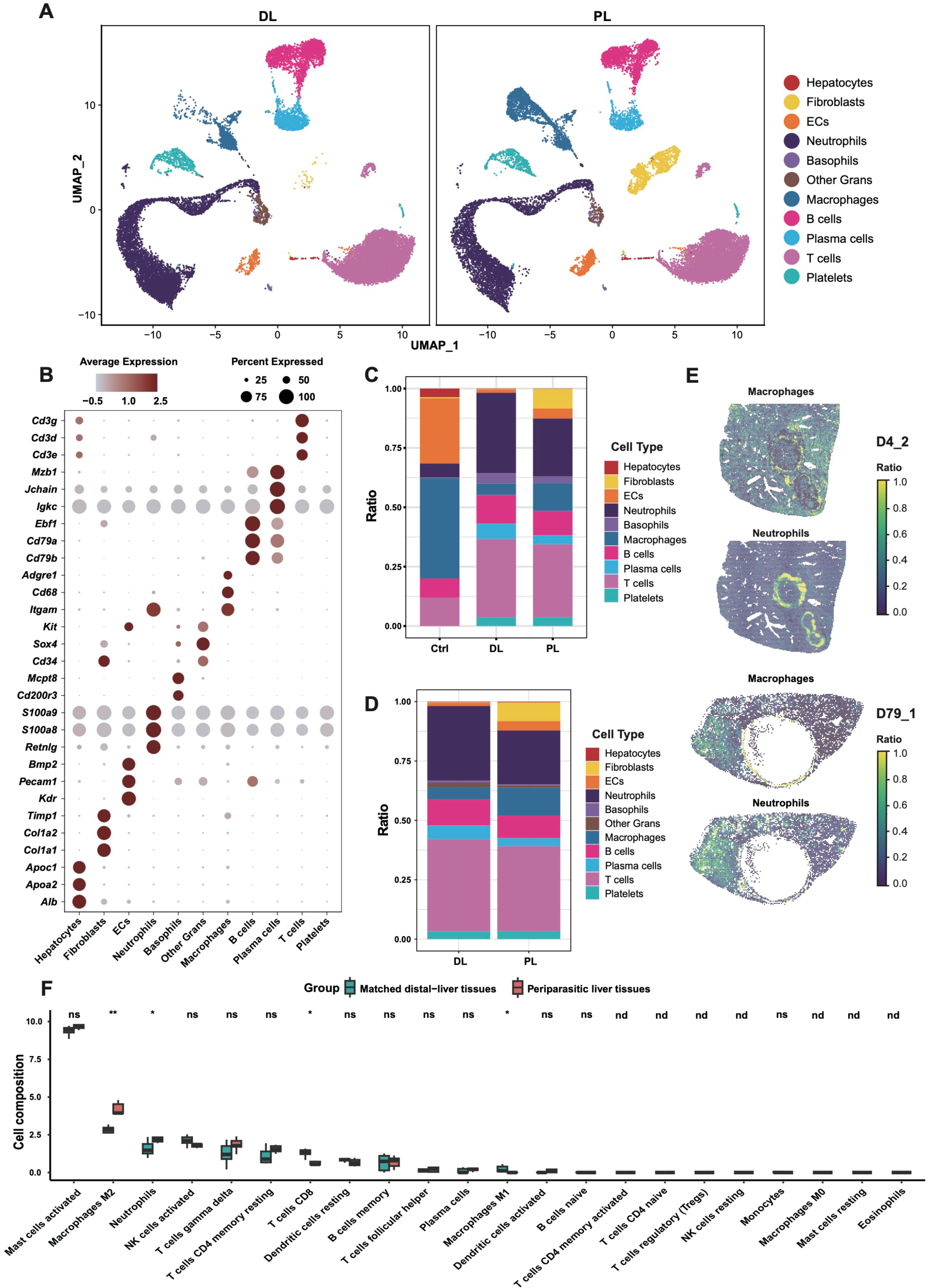
Cellular composition differences between the distal- and peri-lesion region of AE lesions in the liver. (A) UMAP of cells identified from the scRNA-seq data of liver tissues (n = 4) collected from two mice infected by *Echinococcus multilocularis* for 15 months. DL: distal-lesion (left); PL: peri-lesion (right). (B) Expression of cell-type marker genes by the cell types identified from the scRNA-seq data of mouse livers. (C) Comparison of cell type compositions between uninfected liver, DL, and PL groups. (D) Comparison of cell type compositions between DL and PL groups. (E) Deconvolution of neutrophils and macrophages in Stereo-seq data with Tangram using the scRNA-seq data as a reference. A higher ratio indicates a better match of the cell types. (F) Deconvolution of liver bulk RNA-seq data from AE patients. Dataset: GSE124362, n = 6.

The transcriptomic data from mice demonstrated significant infiltrations of neutrophils around AE lesions in the liver (Figures 1B-E, 1G, and 2C). It’s fundamental to confirm if the same phenomena occur in human cases. Based on a public dataset of six AE patients (aged from 33 to 47), GSE124362, we found that the proportion of neutrophils in the periparasitic liver tissues was significantly increased compared with the distal liver tissues (Figure 2F), indicating a pivotal role of neutrophils in the immune response to *E. multilocularis* infection. Unfortunately, although neutrophil counts were elevated in infected individuals ^25^, AE persistently exacerbated, suggesting their inability to eliminate metacestodes.

### Transcriptionally and morphologically heterogeneous neutrophil populations in *Echinococcus multilocularis* infected liver

Visualization of neutrophils showed two types of spatial distributions: aggregation around the AE lesions or sporadically located in the other regions of the infected mouse liver (Figure 2E). To reveal the heterogeneity of neutrophils, we divided the neutrophils identified in the Stereo-seq chips into subclusters and identified two transcriptionally distinct subpopulations, which were designated as *Il1b*^hi^ Neu and *Mpo*^hi^ Neu (Figures 3A and S4A). The *Il1b*^hi^ Neu showed decreased expression of classical neutrophil markers and highly expressed *Fth1, Il1b*, *Ccl3,* and *Ccl4* (Figure 3B). The *Mpo*^hi^ Neu expressed both neutrophil progenitors markers such as *Elane*, *Mpo*, and *Prtn3* and mature markers such as *Camp* and *Ltf* ^26^, suggesting that these cells may be newly recruited to the infection foci and undergoing a phenotype transition. The *Il1b*^hi^ Neu were mainly identified in samples collected on 4 dpi and 8 dpi (Figure 3C). Spatially, the *Il1b*^hi^ Neu were significantly enriched in the AE lesions of these samples (Figure 3D). In contrast, *Mpo*^hi^ Neu tended to be scattered in both infected and non-infected regions (Figures 3C and D). *Mpo*^hi^ Neu were identified in 14 chips including the infected mice and non-infected mice. Morphologically, *Il1b*^hi^ Neu tended to have a multi-lobular or even hyper-segmented nucleus (Figure 3E), which was similar to the neutrophil morphological heterogenicity observed in acute respiratory distress syndrome ^27^, while *Mpo*^hi^ Neu seemed to have a ring-form nucleus, similar to that of blood neutrophils (Figure 3F). IHC staining with IL-1β showed positive signals in neutrophils gathering in the infection foci (Figure S4B), while neutrophils with a ring-form nucleus in the distal region of the lesion were negative (Figure S4C). Meanwhile, staining of myeloperoxidase (encoded by *Mpo*) displayed positive signals in neutrophils with both multi-lobular and ring-form nuclei (Figures S4D and S4E), indicating that the *Mpo*^hi^ Neu might be morphologically dynamic and located both near the infection foci and distal regions. Observation of H&E-stained samples showed that the number of neutrophils gradually decreased and no longer heavily aggregated around the AE lesions as the infection persisted. At the late stage of infection, abundant fibroblasts and inflammatory cells were present around the parasitic microcysts, primarily composed of *Spp1*^+^ MoMFs (Figures S1A and S2E, D79_1), forming a fibrous layer together. This may be related to the immune response transition corresponding to the different stages of AE.

**Figure 3.**
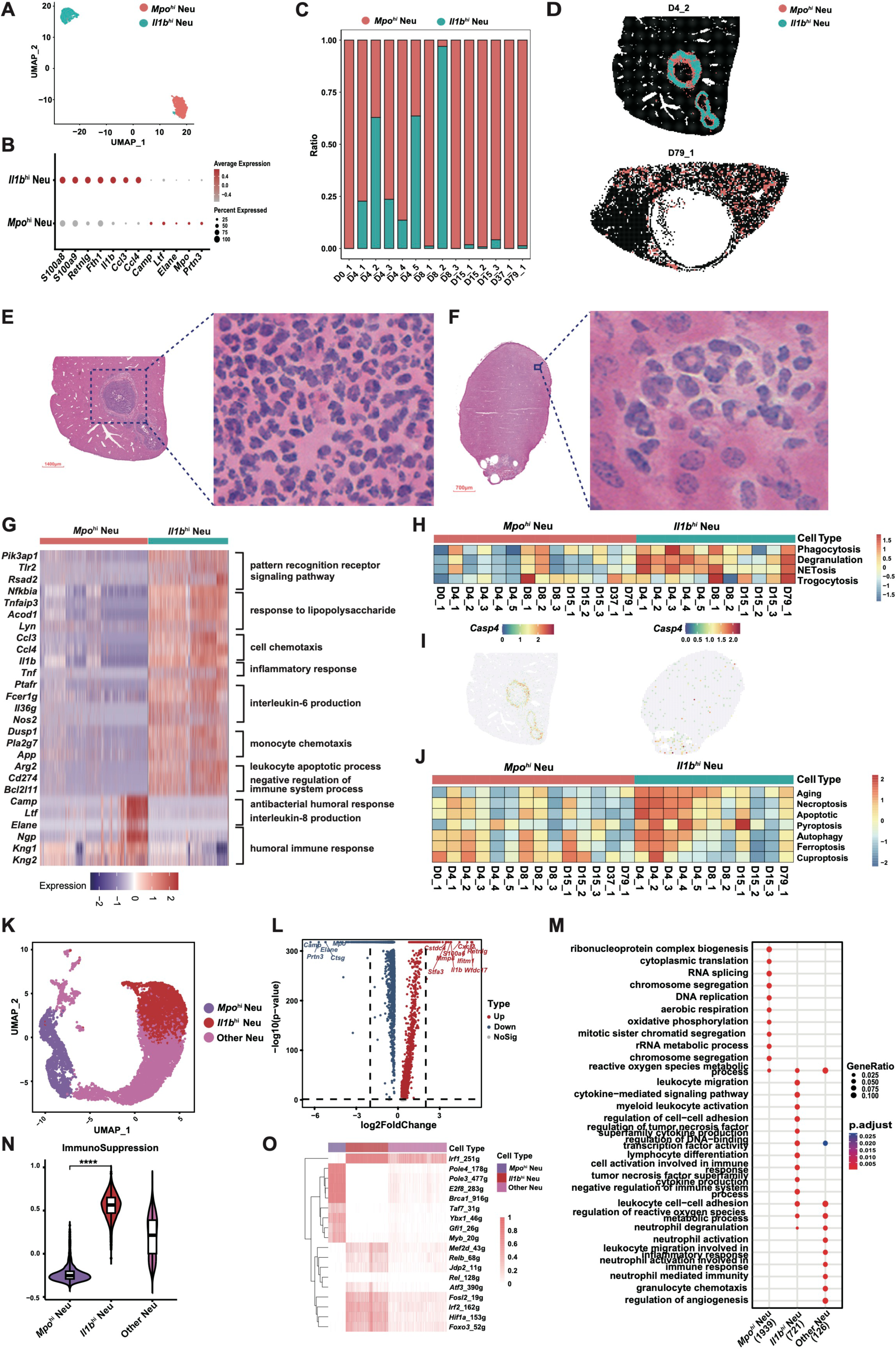
Transcriptional, morphological, and functional heterogeneity of neutrophil subpopulations. (A) UMAP of two neutrophil subpopulations identified from the Stereo-seq data of mouse liver tissues. *Il1b*^hi^ Neu, neutrophils highly expressing *Il1b*; *Mpo^hi^* Neu, neutrophils highly expressing *Mpo*. (B) Dot plot showing the expression of selected DEGs in *Il1b*^hi^ Neu and *Mpo^hi^* Neu. (C) Proportions of *Il1b*^hi^ Neu and *Mpo^hi^* Neu in mouse liver samples collected at different timepoints based on Stereo-seq data. (D) Spatial distribution of *Il1b*^hi^ Neu and Mpo^hi^ Neu in mouse livers infected with *Echinococcus multilocularis.* (E) Neutrophils aggerated at the AE lesion mostly had a multi-lobular nucleus. (F) Neutrophils dispersed diffusely in the infected liver had a ring-form nucleus. (G) Heatmap showing the expression levels of representative genes associated with multiple biological functions for *Mpo^hi^* Neu and *Il1b*^hi^ Neu, standardized by Z-score. (H) Heatmap showing the GSVA scores for pathogen-killing pathways of *Mpo^hi^* Neu and *Il1b*^hi^ Neu based on Stereo-seq data. (I) Spatial expression pattern of *Casp4*, a gene related to NETosis, in sample D4_2 (left) and D8_1 (right). (J) Heatmap showing the GSVA scores for aging and cell death pathways of *Mpo^hi^* Neu and *Il1b*^hi^ Neu based on Stereo-seq data. (K) UMAP of three neutrophil subpopulations identified from the scRNA-seq data of mouse liver tissues. (L) Volcano plot showing the differentially expressed genes between *Mpo^hi^* Neu and *Il1b*^hi^ Neu. Red and blue dots indicate genes significantly upregulated and downregulated in *Il1b*^hi^ Neu. (M) GO enrichment of genes differentially expressed between the three neutrophil subpopulations. (N) Violin plot showing the gene signature scores of the immune suppression pathway for the three neutrophil subpopulations identified from scRNA-seq data. (O) Differentially expressed regulatory genes of the three neutrophil subgroups identified from scRNA-seq data.

### Functional heterogeneity of neutrophils in *Echinococcus multilocularis* infected mouse liver

To further explore the functional differences between the two neutrophil subpopulations, we performed Gene Ontology (GO) enrichment analysis to characterize the enrichment of specific pathways of the two neutrophil populations (Figure 3G). Compared with *Mpo^hi^* Neu, the *Il1b*^hi^ Neu displayed enhanced pattern-recognition receptor (PRR)-mediated response to pathogens (*Tlr2*) and response to lipopolysaccharide (*Tnfaip3*, *Acod1*) ^28^, both were vital pathogen recognition pathways against parasites. Moreover, gene signature analysis indicated that *Il1b*^hi^ Neu had the highest PRR activity among the immune cells, followed by *Mpo^hi^* Neu and *Spp1*^+^ MoMFs (Figure S5A and S5B). In addition, the *Il1b*^hi^ Neu exhibited higher expression of genes primarily associated with chemotaxis (*Ccl3*, *Ccl4*), pro-inflammatory cytokines (*Il1b*, *Tnf*), interleukin-6 (*IL-6*) production (*Ptafr*, *Fcer1g*, *Il36g*, and *Nos2*), monocyte chemotaxis/monocyte chemotactic protein-1 (*MCP-1*) production (*Ccl4, Ccl3, Dusp1, Pla2g7,* and *App*), negative regulation of the immune system (*Arg2*, *Cd274*, and *Dusp1*), and apoptotic process (*Arg2*, *Cd274*, and *Bcl2l11*) (Figure 3G). Pathways such as cell chemotaxis and inflammatory response were associated with the pro-inflammatory role of *Il1b*^hi^ Neu and were consistent with the gene signatures of classic neutrophil properties. IL-6 and MCP-1 may promote the recruitment of monocytes, increasing the mononuclear-cell infiltration to the AE lesion ^29,30^. Interestingly, genes related to the negative regulation of the immune system (*Arg2, Cd274*) were also enriched in *Il1b*^hi^ Neu (Figure 3G), indicating the anti-inflammatory feedback of these cells. The *Il1b*^hi^ Neu, with higher apoptotic signatures, might be more prone to be engulfed by phagocytes ^31^. Comparatively, the *Mpo^hi^* Neu displayed a gene expression signature suggestive of innate antibacterial immunity (*Camp*, *Ltf*, *Elane*) and defense response to pathogens (*Ngp*) (Figure 3G). These *Mpo^hi^* Neu may be newly recruited to the infection tissues, as they displayed both immature and mature neutrophil signatures, whether they would function effectively to control the parasites remains unclear.

Neutrophils can kill extracellular pathogens by phagocytosis, secreting granules with cytotoxic enzymes (degranulation), producing neutrophil extracellular traps (NETs, NETosis), or devouring the content of another organism bit by bit (trogocytosis) ^32,33^. We estimated the GSVA scores of several processes including phagocytosis, degranulation, NETosis, and trogocytosis based on the Stereo-seq data. Results showed the *Il1b*^hi^ Neu displayed higher activities in phagocytosis, degranulation, and NETosis than *Mpo^hi^* Neu (Figure 3H), especially in samples collected on 4 dpi, indicating a stronger parasite-killing capability of *Il1b*^hi^ Neu at the early infection stage. *Casp4*, a gene related to the formation of NETs ^34,35^, was upregulated in *Il1b*^hi^ Neu near the infection foci (Figure 3I). It is reported that trogocytosis was involved in *Trichomonas vaginalis* killing by neutrophils ^33^, but the expression levels of the associated genes varied greatly among our samples (Figure 3H). Notably, *Mpo^hi^* Neu, rather than *Il1b*^hi^ Neu, highly expressed two proteinases (*Elane* and *Prtn3*) in the trogocytosis process (Figure 3B) ^33^, introducing bias into our GSVA scoring (Figure 3H). Thus, we were unsure about the presence of trogocytosis in neutrophils during *E. multilocularis* infection.

The death of the aged and apoptotic neutrophils at the Inflammatory site may be involved in the negative regulation of immune response against infection ^36,37^. We evaluated the gene signature expression levels for cell aging and death pathways based on the Stereo-seq data. The *Il1b*^hi^ Neu exhibited higher scores of aging, necroptosis, apoptosis, pyroptosis, and autophagy compared with *Mpo*^hi^ Neu (Figure 3J). Interestingly, the *Mpo*^hi^ Neu seemed to have higher activity in the cuproptosis pathway (Figure 3J).

To further validate our findings from spatial data, we divided the neutrophils identified in the mouse scRNA-seq data into three subclusters, including the *Mpo*^hi^ Neu, *Il1b*^hi^ Neu, and the others (Figure 3K). The differentially expressed genes between *Mpo*^hi^ Neu and *Il1b*^hi^ Neu from the scRNA-seq data were consistent with those identified from Stereo-seq data (Figures 3L and S5C). Consistently, *Il1b*^hi^ Neu upregulated pathways including leukocyte activation, adhesion, cytokine production, degranulation, and negative regulation of the immune response (Figure 3M). NETosis activity was especially pronounced in *Il1b*^hi^ Neu, both in the DL and PL groups (Figure S5D). Higher signals of aging and death pathways were also observed in *Il1b*^hi^ Neu compared with *Mpo*^hi^ Neu and the other Neu (Figure S5E). Moreover, the *Il1b*^hi^ Neu in the peri-lesion region exhibited a higher potential for aging and death (apoptosis and ferroptosis) than in the distal-lesion region (Figure S5E). Interestingly, the necroptosis activity was higher in the three groups of neutrophils in the distal-lesion region than in the peri-lesion region (Figure S5E). Gene signature analysis revealed the highest activity of immune suppression in *Il1b*^hi^ Neu among the three neutrophil subclusters (Figure 3N). We further used SCENIC to explore the regulatory factors associated with the differential functions of the two neutrophil subpopulations ^38^. *Irf1*, *Irf2*, *Hif1a*, *Foxo3*, and *Fosl2* were closely related to the functional activation of *Il1b*^hi^ Neu, while *Brca1*, *Pole3*, *E2f8*, and *Pole4* were essential to the function of *Mpo*^hi^ Neu (Figure 3O).

Overall, our evidence supports the involvement of functionally heterogeneous neutrophil populations in the parasite control and immune regulation process in livers infected with *E. multilocularis*.

### Cell-cell communications between neutrophils and the other cell types

Cell-cell communication analysis based on the Stereo-seq chips revealed frequent interactions among the different cell types in the AE lesions (Figure S6A). To identify reliable ligand-receptor pairs, we used Stereo-seq data (Figures 4A and 4B) to find interaction pairs that are spatially achievable whereas the scRNA-seq data (Figures 4C and 4D) was used to confirm the most probable cell type origin of each ligand/receptor gene. As the major parenchymal cell type in the liver, hepatocytes are inevitably affected by parasite infection and may release signals to recruit immune cells. Specifically, the hepatocytes may recruit neutrophils through the *Saa2*-*Fpr2* axis (Figures 4A and 4C), consistent with a previous study ^39^. At the same time, a unique high expression of *Saa2* was observed in hepatocytes instead of the other cell types in our scRNA-seq data (Figure 4D).

**Figure 4.**
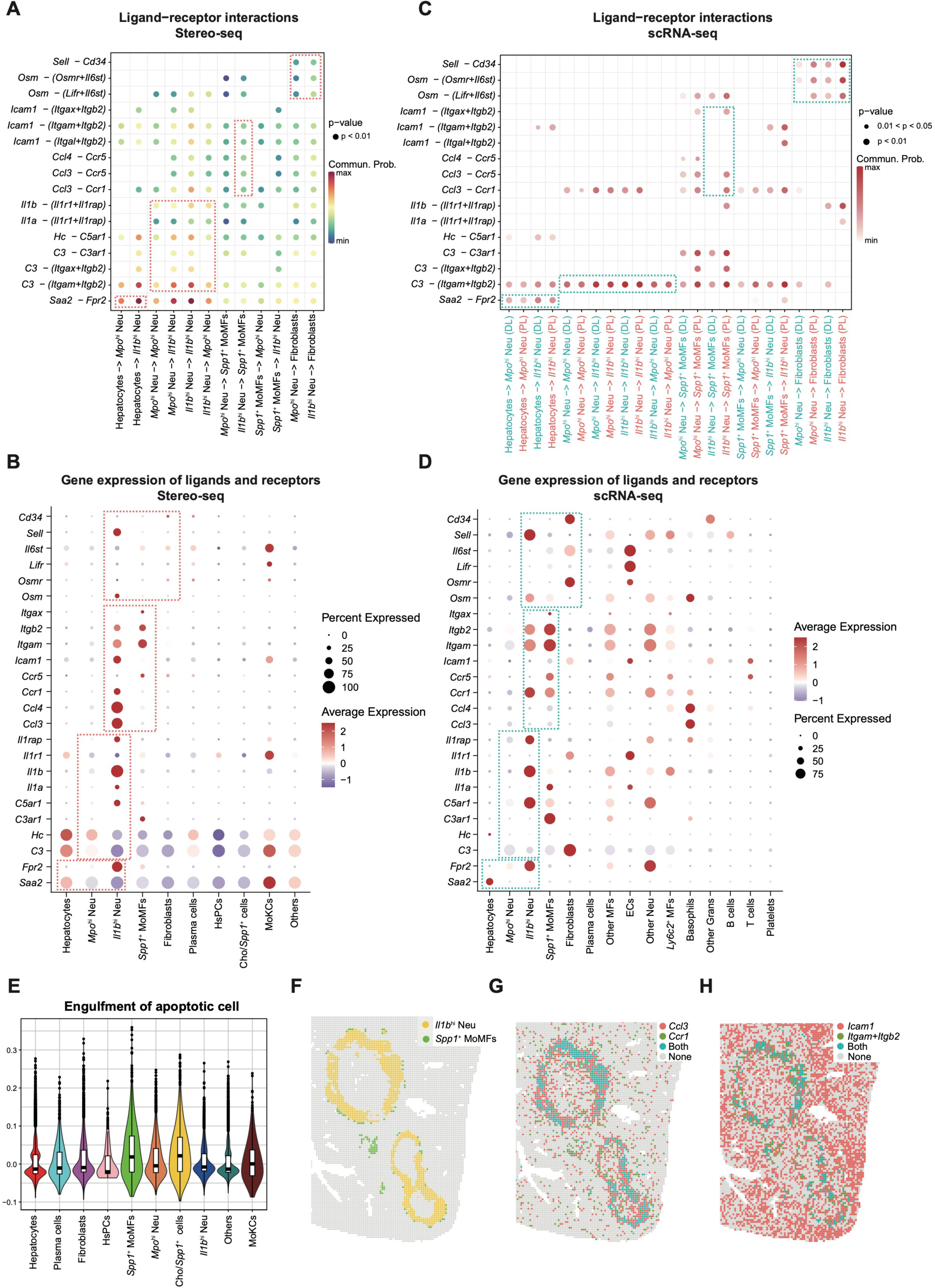
Neutrophil-associated cell-cell interactions in the AE lesions. (A) Bubble plot showing the top ligand-receptor pairs between neutrophils, hepatocytes, *Spp1^+^* MoMFs, and fibroblasts identified by CellChat based on the Stereo-seq data of D4_2. (B) Dotplot showing the expression levels of genes shown in (A) based on cells identified from 14 Stereo-seq chips. (C) Bubble plot showing the significant ligand-receptor pairs shown in (A) between *Il1b*^hi^ Neu and the other cell types (including *Mpo*^hi^ Neu, *Spp1^+^* MoMFs, hepatocytes, and fibroblasts) in the DL and PL groups of the scRNA-seq data. (D) Dotplot showing the expression levels of genes shown in (C) based on cells identified from the scRNA-seq data. (E) Violin plot showing the gene signature scores of engulfment of apoptotic cells for different cell types identified in 14 Stereo-seq chips. (F) Distribution of *Il1b*^hi^ Neu (yellow dots) and *Spp1^+^* MoMFs (green dots) in Sample D4_2. (G) Spatial distribution of *Ccl3*-*Ccr1* ligand-receptor pairs in Sample D4_2. (H) Spatial distribution of *Icam1* - (*Itgam*+*Itgb2*) ligand-receptor pairs in Sample D4_2.

Besides the leukocyte recruiting ability of the injured hepatocytes, *Mpo*^hi^ Neu and *Il1b*^hi^ Neu may further stimulate the migration and aggregation of neutrophils to the infected tissues through ligand-receptor pairs such as *Il1b* - (*Il1r1*+*Il1rap*), *Il1a -* (*Il1r1*+*Il1rap*), *C3*-*C3ar1,* and *C3* - (*Itgam*+*Itgb2*) (Figures 4A and 4C). Furthermore, the *Il1b*^hi^ Neu may recruit *Spp1*^+^ MoMFs through *Ccl3*-*Ccr1*, *Ccl3*-*Ccr5,* or *Ccl4*-*Ccr5* interaction pairs (Figures 4A and 4C), which can be supported by the gene expression profiles in Stereo-seq (Figure 4B) and scRNA-seq data (Figure 4D). It is reported that NETs/NETosis neutrophils can be engulfed by macrophages, called efferocytosis ^40^. In our data, *Icam1* on *Il1b*^hi^ Neu and *Itgam*/*Itgb2* on *Spp1*^+^ MoMFs may be associated with the phagocytosis of local apoptotic *Il1b*^hi^ Neu (Figures 4A and 4C). Besides, the *Spp1*^+^ MoMFs also exhibited higher scores of engulfment of apoptotic cells (Figure 4E). The gene expression of ligands and receptors associated with recruitment and phagocytosis were upregulated in *Il1b*^hi^ Neu and *Spp1*^+^ MoMFs, respectively, as revealed by the Stereo-seq (Figure 4B) and scRNA-seq data (Figure 4D). In addition, these ligand-receptor pairs showed a significant colocalization in Sample D4_2, such as *Ccl3*-*Ccr1* and *Icam1* - (*Itgam*+*Itgb2*) (Figures 4F-4H), supporting the recruitment and phagocytosis between *Il1b*^hi^ Neu and *Spp1*^+^ MoMFs. The neutrophils also interacted intensively with fibroblasts through ligand-receptor pairs such as *Osm* - (*Osmr*+*Il6st*) or *Osm* - (*Lifr*+*Il6st*), as well as the *Sell*-*Cd34* (Figures 4A-4D). NicheNet analysis based on the scRNA-seq data showed that *Il1b*^hi^ Neu upregulated the activity of ligands including *Tnf*, *Il1b*, and *Osm* (Figure S6B), and the functional consequences of their corresponding targets in the fibroblasts may be linked with pro-fibrotic and pro-angiogenesis activities (Figure S6C).

### *Spp1^+^* MoMFs displayed two-sided functions

While neutrophil swarming was evident in the AE lesions, macrophages were also recruited to the infection foci soon after infection. The macrophages surrounded closely around the PSC, possibly trying to engulf the pathogen through phagocytosis. Clustering of macrophages in the mouse liver scRNA-seq dataset revealed four major subgroups, including *Spp1*^+^ MoMFs, MoKCs, *Ly6c*^+^ macrophages (*Ly6c*^+^ MFs), and the other macrophages (other MFs) (Figures 5A and 5B). Analysis of gene signatures showed that the *Spp1*^+^ MoMFs exhibited an M2 (anti-inflammatory) phenotype and may be involved in the angiogenesis process (Figure 5C). These *Spp1*^+^ MoMFs were mostly present in the peri-lesion region (Figure 5D) and were located near the *E. multilocularis* microcysts (Figures 1B and S1A). Our spatial data revealed the appearance of *Spp1*^+^ MoMFs soon after infection, indicating the continuous function of these cells through the early-to-late infection stages. On 4 dpi, the *Spp1*^+^ MoMFs closely attached the PSCs to kill the parasites (Figure 5E, D4_2). When the PSCs developed into microcysts, they continued to gather outside the microcysts (Figure 5E, D79_1). The *Spp1*^+^ MoMFs displayed a pro-inflammatory function (M1 phenotype) from 4 to 79 dpi (Figure 5F). However, their anti-inflammatory function (M2 phenotype) also seemed to increase after one month post infection (Figure 5G). Meanwhile, the *Spp1*^+^ MoMFs showed an enhanced antigen recognition and presentation activity (Figure 5H), and a likely stable phagocytosis activity since 4 dpi (Figure 5I). Intriguingly, we didn’t identify an obvious increase in the gene signature scores for the angiogenesis pathway in *Spp1*^+^ MoMFs during AE progression (Figure 5J). Even so, *Spp1*^+^ MoMFs exhibited a relatively higher signature score for angiogenesis than most of the other cell types, both in our scRNA-seq (Figure S7A) and Stereo-seq data (Figure S7B). Our scRNA-seq data also revealed a high expression level of *Vegfa* in the *Spp1*^+^ MoMFs, which is a critical gene promoting angiogenesis (Figure S7C). We further used SCENIC to explore the regulatory genes associated with the unique function of *Spp1*^+^ MoMFs. Results showed that *Etv1*, *Etv5*, *Bhlhe40, Mitf,* and *Maf* were the most specific regulons in *Spp1*^+^ MoMFs compared with the other macrophages (Figure 5K). Our data suggested a two-sided role of *Spp1*^+^ MoMFs in AE, that they were involved in parasite-killing since the early infection stage but were also closely associated with pathological characteristics in the late infection stage, including immunosuppression and angiogenesis.

**Figure 5.**
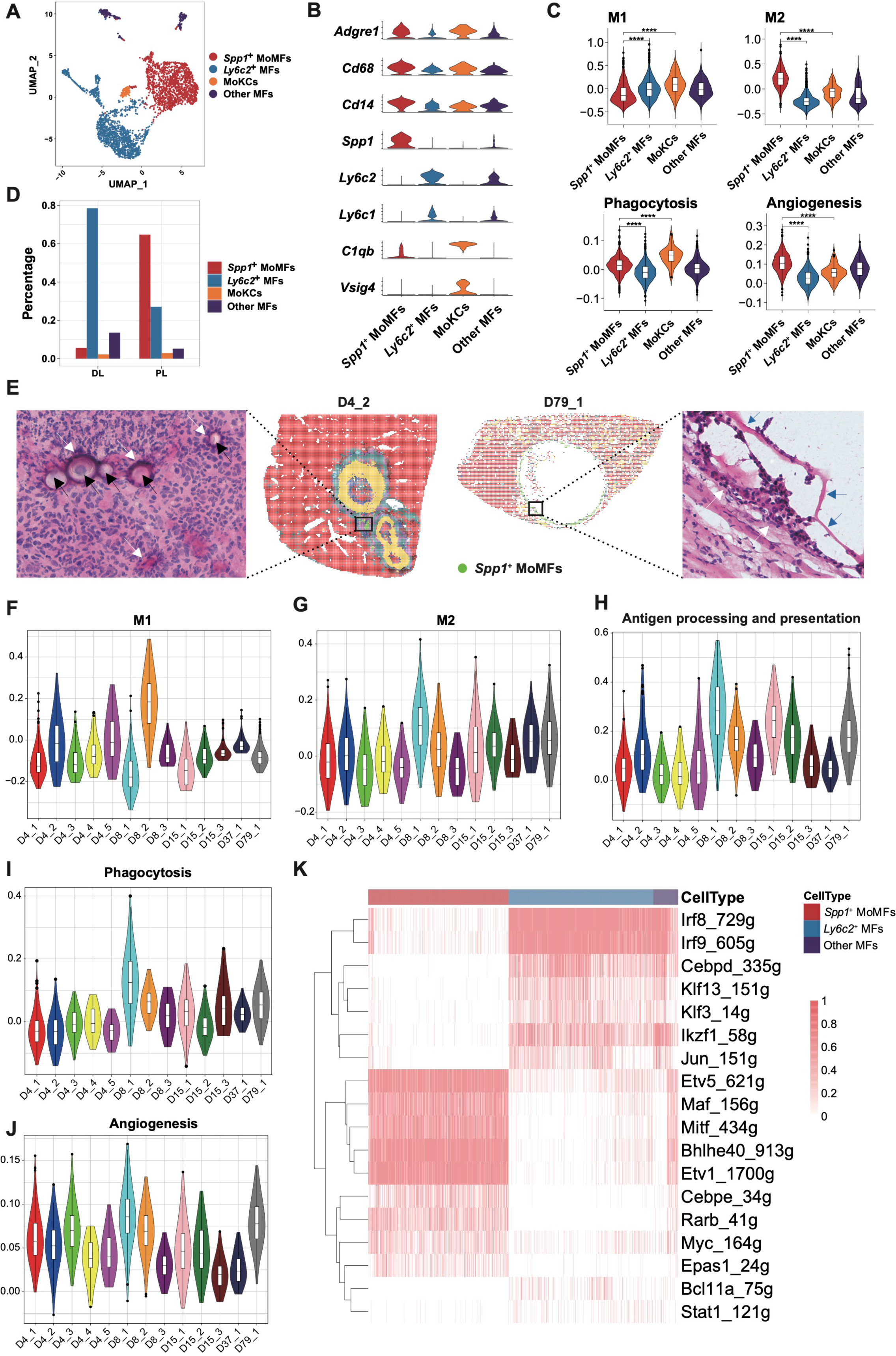
Characterization and functional analysis of *Spp1*^+^ MoMFs. (A) Clustering of the macrophage subpopulations identified from the scRNA-seq data of liver tissues (n = 4) collected from two mice infected by *Echinococcus multilocularis* for 15 months. (B) Expression of marker genes in macrophage subpopulations. (C) Gene signature scores of M1 phenotype, M2 phenotype, phagocytosis, and angiogenesis pathways for the four macrophage subpopulations. (D) Distribution of the four macrophage subpopulations in the DL and PL regions. (E) Stereo-seq chips and H&E staining images showing the spatial distribution of *Spp1^+^* MoMFs in Sample D4_2 and D79_1. Black arrows indicate PSCs. Blue arrows indicate the germinal layer inside the microcyte of *Echinococcus multilocularis.* White arrows indicate *Spp1^+^* MoMFs surrounding the PSC or microcyst. Gene signature scores of M1 phenotype (F), M2 phenotype (G), antigen processing and presentation (H), phagocytosis (I), and angiogenesis (J) pathways for *Spp1^+^* MoMFs identified in Stereo-seq samples collected at 4, 8, 15, 37, and 79 dpi. (K) Differentially expressed regulatory genes of the macrophage subpopulations identified from scRNA-seq data.

### Interactions between *Spp1^+^* MoMFs and fibroblasts may promote fibrosi

Spatially, the *Spp1*+ MoMFs were highly colocalized with fibroblasts, with a correlation coefficient of 0.62 in sample D79_1 (Figure 6A). Cell-cell interaction analysis of scRNA-seq data revealed higher interaction intensity of *Spp1*^+^ MoMFs and fibroblasts in the peri-lesion group than in the distal-lesion group (Figure 6B). The *Spp1*^+^ MoMFs upregulated the activity of *Tgfb1*, *Spp1*, and *Mmp14* (Figures 6C and S8A). *Tgfb1* bound to *Tgfbr1*, *Tgfbr2*, and *Tgfbr3*, which were highly expressed by fibroblasts. *Spp1* interacted with *Itga5*, *ItgaV*, *Itgb1,* and *Itgb5* that were expressed by fibroblasts. The receptor of *Mmp14*, *Cd44,* was also upregulated by fibroblasts (Figure S8A). The simultaneous upregulation of ligands and corresponding targets on the two cell types in shared tissue regions was further confirmed by spatial data (Figures 6D and 6E). Moreover, collagen (*Col1a1*, *Col4a1*, and *Col5a1*) and matrix metalloproteinase (*Timp1*, *Mmp2*) were highly expressed by fibroblasts (Figure 6E). These targets played important roles in pathways associated with extracellular matrix (ECM) remodeling, cell proliferation, and angiogenesis (Figure S8B), which were closely related to the development of liver fibrosis. To confirm the proliferative potential of fibroblasts in AE, we calculated the proliferation gene signature scores for cells in the AE lesions. Results showed that the proliferation scores of fibroblasts were higher in the peri-lesion region than those in distal-lesion region, based on scRNA-seq data (Figure 6F, left). Our Stereo-seq data also showed increased cell proliferation potential of fibroblasts in the infected foci (Figures 6F and 6G). Furthermore, the spatial expression of the fibroblast marker gene set was highly correlated with the cell proliferation gene set in multiple samples (Figure S9A), with a correlation coefficient of 0.39 in sample D4_2 (Figure 6H). Notably, *Spp1*^+^ MoMFs may promote the growth of fibroblasts through the interactions of growth factor associated ligands and receptors including *Pdgfa*-*Pdgfra*/*Pdgfrb* and *Pdgfc*-*Pdgfra,* as shown in our Stereo-seq data and scRNA-seq data (Figure 6I). Intriguingly, we also discovered a high expression of *Cd34* in fibroblasts (Figures 2B and S8A), which was associated with microvascular formation in AE lesions ^41,42^, indicating the potential role of fibroblasts in promoting vessel development. Overall, our evidence indicated that *Spp1*^+^ MoMFs and fibroblasts may interact closely to facilitate fibrosis and vasculature development in the AE lesions.

**Figure 6.**
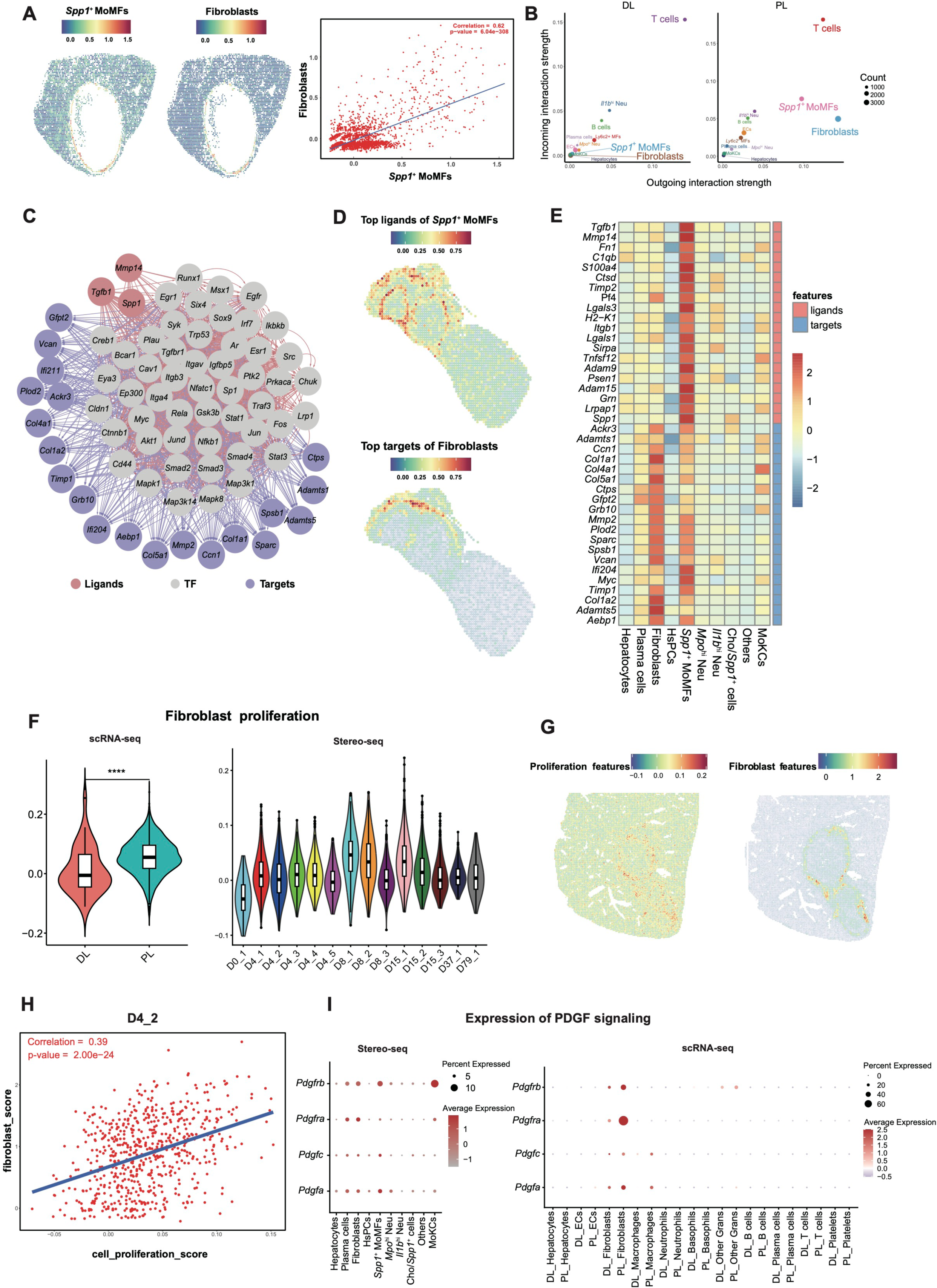
Cell-cell interactions between *Spp1*^+^ MoMFs and fibroblasts. (A) Spatial distribution patterns and correlation analysis of *Spp1^+^* MoMFs and fibroblasts indicated colocalization of the two cell types. (B) Cell-cell communication strength estimated by Cellchat. *Spp1^+^* MoMFs and fibroblasts showed higher interaction intensities in the PL group (right) than in the DL group (left). (C) Regulation network of three active ligands expressed by *Spp1*^+^ MoMFs (*Tgfb1, Spp1,* and *Mmp14*). The analysis was carried out with NicheNet using scRNA-seq data. (D) Spatial presentation of the signature scores for selected ligands on *Spp1*^+^ MoMFs (upper) and the associated targets on fibroblasts (bottom) in sample D8_2. (E) Relative expression of 20 ligands on *Spp1*^+^ MoMFs and 20 associated targets on fibroblasts in spatial clusters in Sample D8_2. (F) Violin plots showing the proliferation scores of fibroblasts in DL and PL groups of scRNA-seq (left) and in Stereo-seq samples collected at different timepoints (right). (G) Spatial presentation of the signature scores for cell proliferation (left) and fibroblast marker gene set (right) in sample D4_2. (H) Correlation analysis of cell proliferation and the fibroblasts signature scores indicated the proliferation of fibroblasts in Sample D4_2. (I) Dotplots showing the expression of genes associated with the PDGF signaling pathway in Stereo-seq (left) and scRNA-seq data (right).

### Functional shift of innate immune response during AE progression

Our results indicated that the host innate immune response may shift from a pro-inflammatory dominant phenotype to an anti-inflammatory dominant phenotype during AE progression. We hypothesize that the hosts adopt an “active killing” strategy at the early stage of infection, which involves the aggregation of immune cells such as *Spp1*^+^ MoMFs and neutrophils to kill the parasite, and then change into a “negative segregation” strategy at the late stage of infection, which involves fibrosis formation surrounding the *E. multilocularis* microcysts trying to contain pathogen growth (Figure 7). When the PSCs enter the liver, they are small in size. *Spp1*^+^ MoMFs attach to the surface of PSCs to destroy their integrity, leading to death. At the same time, a large amount of the *Il1b*^hi^ Neu will swarm around the growing metacestodes and kill them through pathways such as NETosis and phagocytosis. More immune cells, including neutrophils and macrophages, will be continuously recruited to the infection foci through cytokines and chemokines expressed by injured hepatocytes and pro-inflammatory immune cells. Most PSCs will be eliminated during the early infection stage. However, as the remaining microcysts grow larger and larger, it becomes harder for the immune cells to efficiently clear the parasites. Moreover, the immune system would gradually reach the immune checkpoint and enhance the expression of immune suppression factors. While the continuous growth and metastasis of the larvae keep stimulating the host immune system, resulting in sustained recruitment of new immune cells to the infected organ, the parasite-killing capability of the immune cells may be hindered by the immune suppressive signals released from the immune cells arrived earlier at the infection foci (eg. *Cd274* expressed by *Il1b*^hi^ Neu, Figure 3G). The M2 phenotype of *Spp1*^+^ MoMFs has been eminent since their appearance (4 dpi), indicating the early expression of anti-inflammatory signals in the AE lesion. Moreover, these *Spp1*^+^ MoMFs can interact with the fibroblasts to stimulate fibroblast proliferation and remodel ECM, which will promote the progression of fibrosis. The formation of fibrotic tissues may help restrict the growth of metacestodes and prevent the spread of parasitic cells into other regions. This “negative segregation” strategy can protect the host organs from parasitic invasion to some extent. On the other hand, while neutrophils continuously migrate into the infected liver, the fibrotic tissues surrounding the AE lesions may hinder their adhesion to microcytes to break down the parasitic components. The *Spp1*^+^ MoMFs become the dominant immune cells surrounding the microcysts, but their parasite-killing capabilities are insufficient regarding the big sizes of the microcytes and the increased thickness of the cystic walls. Moreover, *Spp1*^+^ MoMFs may facilitate angiogenesis in the infection foci, which can provide nutrients for the pathogen, thus accelerating metastasis and exacerbating disease severity.

**Figure 7.**
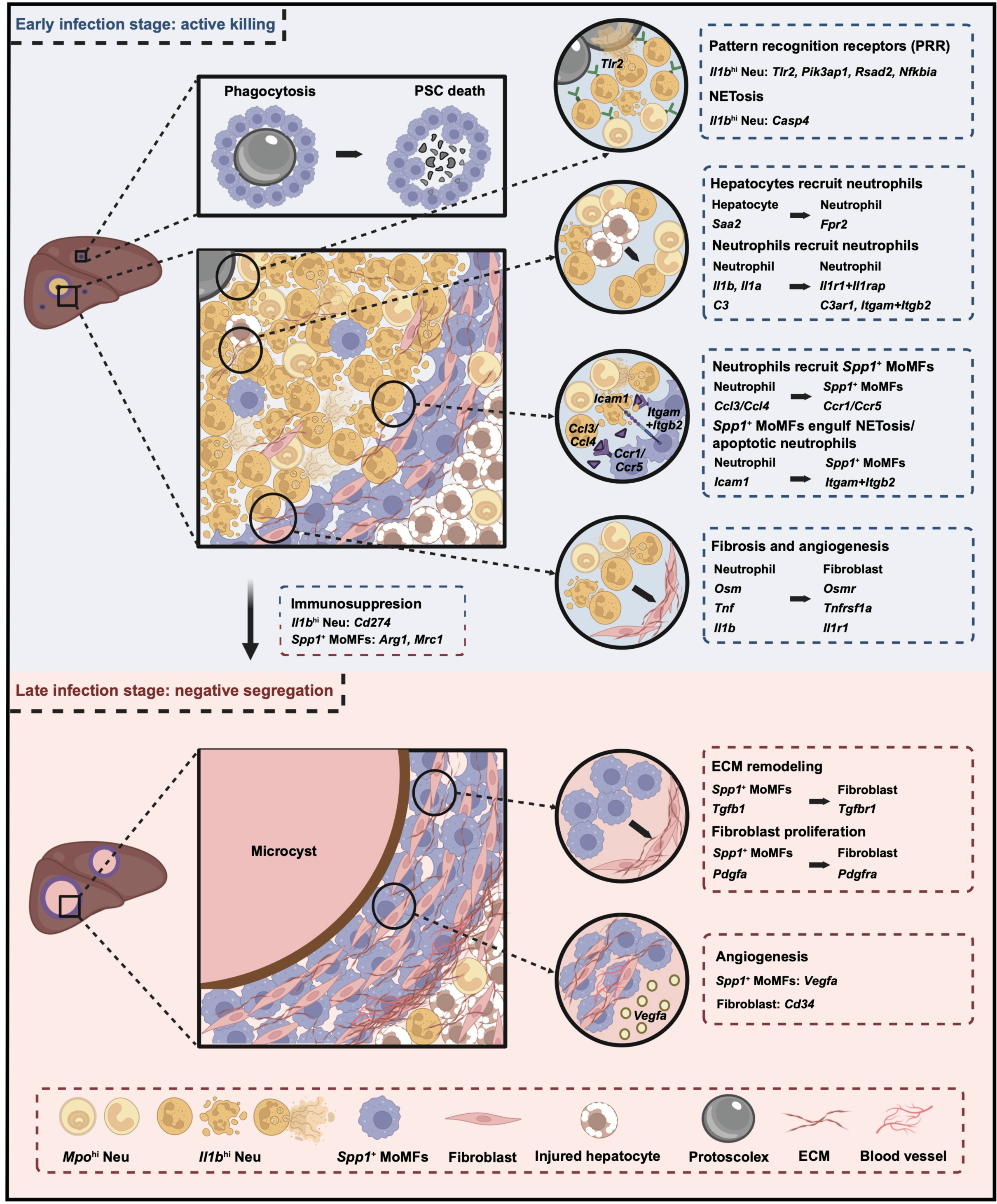
Schematic graph showing the functional involvements of neutrophils, macrophages, and fibroblasts during AE progression. During the early infection stage, i.e., *Echinococcus multilocularis* in the form of protoscoleces (PSCs) or newly formed microcysts, the host immune cells are recruited to the infected foci to kill the parasites actively (“active killing). Specifically, the *Spp1*^+^ MoMFs attach to and phagocytise the PSCs. Meanwhile, the injured hepatocytes recruit a large amount of *Il1b*^hi^ Neu that highly express pattern-recognition receptor (PRR) to the lesion site, which recognize and clear the parasite through pathways such as NETosis. During this process, more neutrophils and *Spp1*^+^ MoMFs are recruited to the lesion by cytokines released by *Il1b*^hi^ Neu. The *Spp1*^+^ MoMFs also help engulf the apoptotic *Il1b*^hi^ Neu. Meanwhile, the interactions between *Il1b*^hi^ Neu and fibroblasts may facilitate fibrosis and angiogenesis. When the infection sustains, the immunosuppressive signals expressed by *Il1b*^hi^ Neu and *Spp1*^+^ MoMFs may trigger the conversion to a “negative segregation” strategy. At this stage, the microcytes are large and their invasive growth may endanger the nearby tissues. Continuous accumulation of neutrophils and *Spp1*^+^ MoMFs can still be observed in the infected tissues. While *Spp1*^+^ MoMFs can attach to the outer surface of the microcysts, *Il1b*^hi^ Neu fail to migrate into the lesion to kill the metacestodes. Although *Spp1*^+^ MoMFs may function to engulf parasitic components, their interactions with fibroblasts may enhance the proliferation of fibroblasts, extracellular matrix (ECM) remodelling, and angiogenesis. The fibrotic structure surrounding the microcytes may constrain the growth of *Echinococcus multilocularis* and prevent the spread of metacestodes into nearby tissues, but this fibrotic barrier may also hinder the infiltration of immune cells such as neutrophils into the lesion center to eliminate the parasite. Therefore, this process is termed as “negative segregation”.

Regarding the changes in host immune response and tissue structure, the pathogen-killing capacities of immune cells are the highest at the early infection stage, or, more specifically, when the parasites take the form of PSCs or small microcysts. The host immune response, combined with anti-parasite drugs such as albendazole will help eliminate metacestodes from the body. However, when the microcysts grow bigger, parasite clearance becomes harder. On one hand, it is difficult for the immune cells to break down the parasites. On the other hand, destroying the integrity of the microcysts will release many exogenous proteins into the host tissues, which may lead to severe allergies. Another risk is the release of parasitic stem cells, which may cause metastasis into other parts of the body. Therefore, for AE patients at the late infection stage, surgical removal of the microcysts without breaking their integrity would be critical to get rid of the pathogen. In case of potential metastasis, simultaneous administration of anti-parasitic drugs and immunotherapy after surgical operation are also recommended to restrict metacestodes growth and rescue the pathogen-killing function of immune cells such as neutrophils.

## DISCUSSION

Immune suppressions are associated with AE deterioration, as demonstrated by the accelerated disease progression in patients with AIDS or other immunosuppressive conditions ^43^. Experimental studies also reveal the upregulation of immune inhibitive signals in the late stage of *E. multilocularis* infection, with a focus on T cells and macrophages ^18,20,44–46^. Recent studies in cancer research have revealed the functional heterogeneity of neutrophils, which could negatively modulate both innate and adaptive immunity ^47,48^. It is reported that the number of neutrophils was increased in AE patients ^25^, yet little is known about its functional role during long-term infection. Using high-resolution spatial transcriptomic technology, we show that the neutrophils in AE lesions were both functionally and morphologically heterogeneous, and the balance between the pro-inflammatory and anti-inflammatory phenotypes of these cells may be critical for the clearance of *E. multilocularis.* Both animal and human data confirmed the increase of neutrophils in infected individuals ^25^, however, the infection were not effectively restrained, indicating the inability of parasite-killing function of these cells in the long run.

Under certain infection circumstances, it is possible to observe the release of both immature and mature granulocytes from the bone marrow simultaneously ^49^. In this study, we have identified a group of partially immature neutrophils, i.e., the *Mpo*^hi^ Neu, indicating abnormal granulocyte release during *E. multilocularis* infection. Myeloid-derived suppressor cells (MDSCs), including neutrophils, were found to be involved in cancer progression and indicative of prognosis and therapeutics ^50^. While we have found that the *Il1b*^hi^ Neu might have acquired the capacity to suppress immune responses during AE, more evidence is needed to understand the differentiation trajectory of these neutrophil subsets.

Neutrophil death plays a critical role in regulating neutrophil-mediated innate immunity, which is mainly anti-inflammatory ^36^. Here, the *Il1b*^hi^ Neu expressed many features of aging and apoptosis, which may negatively regulate host immunity. It was reported that the delay of neutrophil death by GSDMD deficiency augmented host defense against extracellular *Escherichia coli* infection ^37^. However, regarding the sustained increase of neutrophils but inefficient parasite inhibiting effects of these cells in AE patients, delaying neutrophil death may be unpractical in treating *E. multilocularis* infection. It would be more suitable to rescue the pathogen-killing function of neutrophils at the infection foci. The cell type composition changes in the AE lesions from early to late infection stages suggested the intensified recruitment of *Spp1*^+^ MoMFs but also the disappearance of neutrophils outside the microcysts. This may be achieved through local death and clearance of aged neutrophils at the infection foci. In this study, the dying neutrophils may be engulfed by the macrophages through efferocytosis ^51^.

Previous studies of *Echinococcus granulosus* reported an immunosuppressive role of *Spp1*^+^ macrophages in cyst echinococcosis progression, which were expanded at 6 months post infection and may be ineffective in parasite control ^52^. Interestingly, our spatial data showed that the *Spp1*^+^ MoMFs were present at a very early stage after *E. multilocularis* infection (4 dpi) and may have been actively involved in pathogen-killing since the early infection stage. At the same time, they also displayed anti-inflammatory phenotypes and intensively interacted with fibroblasts, participating in ECM remodeling of the AE lesions. It was reported that the *E. multilocularis* antigen can affect the polarization of macrophages and activate the transformation of fibroblasts to induce fibrosis ^46,53^. Our findings also suggest the involvement of *Spp1*^+^ MoMFs in angiogenesis in the AE lesions. It will be interesting to know how this process is triggered by the interaction between parasitic and host factors to facilitate the long-term survival of *E. multilocularis* in the host.

Several pathways may be involved in the clearance of *E. multilocularis* from the liver. Our analysis suggests that NETosis may be one of the parasite-killing mechanisms, but the other pathways can’t be disregarded. Trogocytosis indicates a process in which one cell bites another cell, leading to the transfer of the cellular content of the target cell to the cell initiating the process, which can occur between multiple cell types ^54^. Recent studies show that trogocytosis is involved in the killing of multicellular pathogens, for example, trogocytosis by macrophages kills *Schistosoma japonicum* and trogocytosis by neutrophils kills *Trichomonas vaginalis* ^33,55^. We have tried to identify signals of trogocytosis, however, the associated gene expression levels varied greatly between samples, and we were unable to confirm the occurrence of trogocytosis during *E. multilocularis* infection. Because we were unable to provide evidence of other techniques, whether trogocytosis or other forms of phagocytosis are involved in the parasite killing process remains to be investigated.

In this study, detection of the *E. multilocularis* genes was insufficient, possibly due to the small number of microcysts and the interference of parasitic vesicular fluid in the mRNA release and capture process. Even for the detected parasitic genes, their functions largely remain unknown because of a lack of annotation. It is proposed that the parasite genes can actively regulate the host immune system, yet how the helminthic cells interact with the host cells is unclear. Technical improvements are required to adequately capture both the parasite and host mRNAs so that the cellular and molecular interactions between the two organisms can be fully explored. Identification of the key genes associated with the propagation of *E. multilocularis* in the liver may provide targets to eradicate this parasite from infected tissues.

This study has depicted an unprecedentedly high-resolution cell atlas of AE lesions in mouse liver and elucidated the cellular interaction networks between neutrophils, macrophages, and fibroblasts based on spatial information, revealing their dynamic roles in disease development. Further experimental verification of the molecular interactions and biological consequences would facilitate the discovery of intervention targets for AE. Immunotherapy targeting immune checkpoint blockades such as PD-L1 and TIGIT has produced some promising results in treating AE ^56–58^. Recently, neutrophils are also regarded as a potential target in immunotherapy for cancers ^48^, whether the interventions against neutrophils could benefit AE treatment may be worth exploring.

## METHODS AND MATERIALS

### Animal infection and sample collection

BALB/c mice were purchased from the Experimental Animal Center of Lanzhou Veterinary Research Institute, Chinese Academy of Agricultural Sciences. The protoscoleces of *E. multilocularis* were collected from naturally infected Qinghai vole (*Lasiopodomys fuscus*) in Yushu, Qinghai province, China. The protoscoleces were intraperitoneally inoculated into BALB/c mice in the laboratory to maintain the parasites. Before the infection experiment, protoscoleces were collected from infected mouse liver and examined under an optical microscope to confirm their vitality. After quantification, the innate BALB/c mice were inoculated with 2,000 protoscoleces by hepatic capsule injection or intraperitoneal injection. Mice were kept under standard conditions and euthanized on 4, 8, 15, 37, and 79 dpi to collect liver tissues. The negative control group was not treated and was sacrificed on the day of inoculation of the experimental group. All liver tissues were embedded in pre-cooled OCT (Sakura, USA) shortly after surgery and stored at -80°C for subsequent Stereo-seq experiments.

### Quality control of OCT-embedded samples

For each OCT-embedded sample, 100-200 μm thick sections were cut and used to extract total RNA using the RNeasy Mini Kit (Qiagen, USA). The RNA integrity number (RIN) was checked by a 2100 Bioanalyzer (Agilent, USA) and samples with RIN ≥ 7 were selected for downstream experiments. For each sample, cryosections with a thickness of 10 μm were obtained for hematoxylin and eosin (H&E) (Beyotime Biotechnology, China) to examine the tissue morphology. Samples with appropriate infection foci of *E. multilocularis* and adjacent host tissues were selected for the Stereo-seq experiment.

### Stereo-seq library preparation and sequencing

The spatial transcriptomics experiment was performed using the STOmics gene expression assay kit (BGI-Shenzhen, China) according to the manufacturer’s protocol. The size of the Stereo-seq chips was 1 x 1 cm^2^. The capture spots were 220 nm in diameter and the distance between the centers of two adjacent spots was 500 nm ^22^. The capture probes contained a 25 bp CID barcode, a 10 bp MID, and a 22 bp poly-T to capture mRNAs. In brief, four to six serial cryosections with a thickness of 10 μm were cut from the OCT block containing the mouse liver tissue. One tissue section was placed onto the Stereo-seq chips and quickly incubated at 37°C for 3 minutes. After a 40-minute methanol fixation, the section was subjected to permeabilization by incubating at 37°C for 12 minutes to allow mRNA *in situ* hybridization. Following mRNA capture, reverse transcription (RT) was performed at 42°C for 1 hour. The RT products were subsequently subjected to cDNA release enzyme treatment overnight at 55°C. The released cDNA was purified using DNA clean beads and amplified with PCR mix. The PCR products were used for library construction and finally sequenced with a 50 + 100 bp strategy on an MGI DNBSEQ T series sequencer. One adjacent tissue section was used for H&E staining to facilitate pathological assessment.

### Bulk RNA-seq experiment

When cutting cryosections for the spatial transcriptomics experiment, we collected tissue sections with a thickness of 100-200 μm for a bulk RNA-seq experiment. Total RNA was extracted using the RNeasy Mini Kit (Qiagen, Germany). The RNA concentration was determined by Qubit dsDNA HS Assay kit (Invitrogen, US) using Qubit 4.0 (Invitrogen, US). 1 μg of total RNA was used for bulk RNA sequencing (RNASeq for mRNA 200-400 bp). Sequencing was conducted on the DNBSEQ platform with a paired-end 100 bp strategy and the raw data size for each sample was 20 Gb.

### H&E and IHC staining of mouse liver sections

For liver specimens used for Stereo-seq, a 10-μm-thick consecutive slice was reserved for each sample for H&E staining to identify the parasite infection foci and pathological changes. For the detection of neutrophils, IHC staining was applied to examine the expression of the protein encoded by *Mpo* (Myeloperoxidase, 1:1000, ab208670, Abcam) and *Il1b* (IL-1 beta, 1:100, ab283818, Abcam).

### Single cell isolation, library construction, and sequencing for scRNA-seq

Two mice that had been infected with *E. multilocularis* for 15 months were selected to collect liver specimens. The area within 0.5 cm around the AE lesion was designated as the peri-lesion liver region (peri-lesion, PL), while the area beyond 0.5 cm was designated as the distal lesion liver region (distal-lesion, DL). Each liver was cut into PL and paired DL samples. The tissues were cut into 1 mm^3^ pieces, digested, washed, and resuspended for scRNA-seq library construction using a Chromium Single Cell 3′ Reagent Kit (10× Genomics) as previously described ^59^. The libraries were sequenced on the Illumina NovaSeq 6000 (Illumina, USA) system.

## Bioinformatic analysis

### Processing and analysis of ST data

#### Preliminary processing of Stereo-seq data

The Stereo-seq raw data were automatically processed using the BGI STOmics analytical pipeline (https://cloud.stomics.tech/), including barcode demultiplexing, adapter filtering, mapping to reference genomes, deduplication, and quantification. The reference genome used for this project was a combined genome containing the mouse genome (GRCm39) and the *Echinococcus multilocularis* genome (WormBase release WBPS15). *E. multilocularis g*enes with counts < 10 per chip were defined as background noise and removed. A bin size of 100 (bin100, 100 x 100 spots, i.e., 49.72 x 49.72 µm) was used as the analytical unit, so that each bin100 contains over 1000 mouse gene types for most chips. Data from the tissue-covered area were extracted based on the ssDNA and H&E staining images using the Lasso function of the BGI STOmics website. The generated gem (gene expression matrix) files were analyzed with Seurat v4 ^60^. The chips were normalized using the SCTransform function. Dimension reduction was performed using PCA at a resolution of 0.8 with the first 30 PCs. Unsupervised clustering of bins was performed using UMAP.

#### Detection of Echinococcus multilocularis genes in Stereo-seq chips

We used the PercentageFeatureSet() function in Seurat v4 to calculate the percentage of parasitic genes in each Stereo-seq chip ^60^. All the parasitic gene labels started with EmuJ. The filtering threshold for parasitic genes in each chip was sample-specific and the pathological assessment of H&E staining images of the adjacent tissue section was used as a reference to remove false positive signals.

#### Annotation of bin clusters in Stereo-seq slides

The spatial expression patterns of genes in Stereo-seq slides were conducted with the SpatialFeaturePlot function of Seurat v4 ^60^. We used the merge function provided by Seurat to annotate clusters shared by multiple chips. The harmony algorithm in Harmony R package was used to eliminate the batch effect ^61^. Merged data clustering was performed using PCA at a resolution of 0.5 with the first 15 PCs. Unsupervised clustering of bin100s was performed using UMAP. Cell type probabilities were calculated for each bin100 using factor analysis *via* FindTransferAnchors and TransferData functions in Seurat ^60^. The murine liver reference datasets (https://www.livercellatlas.org/) were used as the single-cell reference to predict the cell composition of each bin100 for all slides. The positive markers for each given marker cell type were identified using the Seurat FindAllMarkers function ^60^. The annotation result was further corrected based on the expression of marker genes. The annotated ST areas were confirmed to be consistent with the pathological assessment and marker gene expression patterns. The H&E images were examined by professional pathologists to determine the tissue types and infection foci.

#### Estimation of cell types and biological activities circling the AE lesions

For each Stereo-seq chip, the center of the AE lesion was determined based on histological assessment based on the H&E staining image of the consecutive tissue section and/or the distribution pattern of *E. multilocularis* genes. The function PercentageFeatureSet in Seurat was used to calculate the percentage of UMIs belonging to *E. multilocularis* for each bin ^60^. The specific filtering threshold values were as follows: 0.05 for D8_2; 0.1 for D4_1, D4_2, D4_3, D4_4, D4_5, D8_3, and D15_1; 0.2 for D8_1 and D15_3; 0.5 for D15_2 and D37_1; 0.8 for D79_1. We expanded the circle from the AE lesion center to distal-lesion regions using 100 μm (two bin100) as a sliding unit. We then calculated the proportions of cell types and performed GSVA for each layer.

#### Functional analysis of neutrophil subpopulations based on Stereo-seq data

The neutrophils identified in Stereo-seq chips were merged and re-clustered into two subpopulations with Seurat. For the pathway enrichment analysis, DEGs expressed by two subpopulations of neutrophils were loaded into R for gene ontology (GO) term enrichment analysis. For neutrophils, gene signature scores of multiple pathways were calculated. Functional signatures for phagocytosis, degranulation (GO:0043312), NETosis, trogocytosis, and cell death-related signatures for neutrophil (necroptosis, apoptotic, pyroptosis, autophagy, ferroptosis, and cuproptosis) were all provided (Table S3). Signatures of neutrophil aging were defined based on literature review ^62^.

#### Co-localization analysis of Spp1^+^ MoMFs and fibroblasts

*Spp1*^+^ MoMF and fibroblast signature (Table S3) scores were calculated based on Stereo-seq data. Each bin100 in clusters located in the lesion region was scored with the AddModuleScore function with default parameters in Seurat. The Pearson correlation of the signature scores of *Spp1*^+^ MoMFs and fibroblasts was calculated with R.

#### Ligand-receptor interaction analysis of Stereo-seq data

We used CellChat (v.2.0.0) to identify the ligand-receptor interaction pairs between cells with space proximity, with the true scale factor for bin100 as follows: spot diameter = 49.72 and spot = 1 ^63^.

### Processing and analysis of scRNA-seq data

#### scRNA-seq data processing

After sample demultiplexing, barcode processing, and single cell 3′ gene counting were performed with Cell Ranger (10x Genomics, v.2.1.1) to acquire a gene-cell matrix using the mouse genome GRCm38 as reference. The matrix data were then filtered with Seurat (v4.3.0) based on the following criteria ^60^: 1) each gene should be present in at least three cells, 2) each cell should contain 500-5,000 genes. For each sample, Ambient RNA and potential doublets were removed using DoubletFinder (v2.0.3) with default settings, respectively. Cells containing over 5,000 genes or had over 10% of their genes belonging to mitochondrial genes were removed.

#### Clustering and annotation of scRNA-seq data

The expression matrices of each sample were integrated with the FindIntegrationAnchors and IntegrateData functions in Seurat (v4.3.0) using the RPCA algorithm ^60^. Downstream analysis such as normalization, log-transformation, highly variable genes identification (2,000 genes excluding mitochondrial and ribosomal genes), dimension reduction, UMAP clustering (dims = 1:30), and differential expression (DEGs) analysis were all conducted with Seurat v4.3.0. After clustering, the cell types were annotated based on reported cell marker genes (Table S2).

#### Functional analysis of scRNA-seq data

Gene ontology (GO) analyses of DEGs in different clusters were performed with clusterProfiler (version: 4.8.2) ^64^. Gene signature scores for different subclusters of macrophages were calculated based on scRNA-seq data. Scores for M1 phenotype, M2 phenotype (Table S3), angiogenesis (GO:0045766), and phagocytosis (GO:0006911) in macrophages were calculated using the AddModuleScore function with default parameters in Seurat. For neutrophils, the gene sets of multiple pathways were as same as those used in Stereo-seq data analysis (Table S3).

#### Ligand-receptor interaction analysis

Ligand-receptor interaction analysis of scRNA seq data was performed using the CellChat (v.1.6.1) R package with default parameters. NicheNet was also used to predict the possible ligand and receptor interactions between *Spp1^+^* MoMFs and fibroblasts, *Il1b^hi^* Neu and fibroblasts based on scRNA-seq data ^65^. The top 67 ligands of *Spp1^+^* MoMFs and the top 440 targets of DEGs of fibroblasts (distal-lesion versus peri-lesion tissues) were extracted for paired ligand-receptor activity analysis. When evaluating the regulatory network of *Spp1^+^* MoMFs on fibroblasts, fibroblasts were considered as receiver cells to check the regulatory potential of *Spp1^+^* MoMFs on fibroblasts. To further explore the consequences of the interaction between *Spp1^+^*MoMFs and fibroblasts, we analyzed the gene set (corresponding targets in fibroblasts of high activity ligands in *Spp1^+^* MoMFs) function in fibroblasts regulated by *Spp1^+^* MoMFs using Metascape ^66^. The interactions between *Il1b^hi^* Neu and fibroblasts were estimated using NicheNet similarly, only that the number of ligands in *Il1b^hi^* Neu and the number of targets in fibroblasts were 36 and 99, respectively.

#### Transcription factor analysis

The transcription factor analysis for neutrophils and macrophages was conducted using SCENIC program (version 1.1.2.2) ^38^ based on scRNA-seq data. We identified transcription factor binding motifs utilizing the gene database from the RcisTarget package (version 1.2.1). The activity of regulons within each cell was scored employing the AUCell package (version 1.4.1). The RSS (Regulon Specificity Score) was calculated based on the Jensen-Shannon divergence (JSD) to assess the specificity of each predicted regulon for different cell types. This involved calculating the JSD between each vector of binary regulon activity overlapping with the assignment of the cells to a specific cell type. The connection specificity index (CSI) for all regulons was determined using the scFunctions package (https://github.com/FloWuenne/scFunctions/).

### Processing and analysis of bulk RNA-seq data

#### Processing of bulk RNA-seq data

Raw FASTQ files were filtered with SOAPnuke (version 2.1.0) and fastp (version 0.20.1) to remove low-quality reads. The reference genome was constructed using the genomes of *Mus musculus* (GRCm39) and *E. multilocularis* (WormBase version WBPS15). The reads were mapped against the reference genomes with STAR (version 2.7.6a), subsequently using salmon (version 1.3.0) for gene expression quantification. 44,908 *Mus musculus* and *E. multilocularis* genes were obtained, generating the raw gene expression count matrix. To facilitate result interpretation, only the following gene types including immunoglobulin variable chain and T-cell receptor genes, lncRNA, protein-coding genes, and tec protein tyrosine kinase genes, were retained in the mouse count matrix, which contained 29,064 genes. The mouse count matrix was TPM normalized, and genes with a TPM score <1 were removed. Finally, the TPM matrix and count matrix for a total of 12,834 mouse genes were generated. The mouse gene matrix was normalized and subjected to principal component analysis (PCA) via the variance Stabilizing Transformation and plotPCA function of DESeq2 (version 1.40.2). The mouse gene identity types were transformed from ENSEMBL to ENTREZID via org.Mm.eg.db (version 3.17.0). The above-mentioned software used default parameters except specially indicated.

#### Deconvolution of bulk RNA-seq data

A signature matrix file was constructed based on our mouse liver scRNA-seq dataset. We then applied utilized CIBERSORTx (https://cibersortx.stanford.edu/) to deconvolute the bulk RNA-seq data of mouse livers obtained in this study for immune cell signals. To test the immune cell composition in the livers of AE patients, we used CIBERSORTx to identify the abundance of immune cells in the GSE124362 dataset using LM22, which was a validated leukocyte gene signature matrix provided by CIBERSORTx, as a signature matrix.

### Statistical analysis and plotting

Statistical analyses, including Student’s t-test, Wilcoxon’s rank-sum test and Wilcoxon signed-rank test, were performed in R 4.0. Asterisks indicate the significance levels of p-values: *, p < 0.05; **, p < 0.01; ***, p < 0.001. The figures were created with ggplot2 in R and BioRender (https://biorender.com/).

### Data and code availability

The data supporting the findings of this study have been deposited into CNSA (CNGB Sequence Archive) of CNGBdb (https://db.cngb.org/cnsa/). The Bulk RNA-seq, scRNA-seq, and Stereo-seq datasets are under the accession numbers of CNP0005294, CNP0005291, and CNP0005297, respectively.

## ETHICAL STATEMENT

This study was reviewed and approved by the Institutional Review Board of Lanzhou Veterinary Research Institute, Chinese Academy of Agricultural Sciences (NO. LVRI-IRB 201911-T1) and Beijing Genomics Institute, Shenzhen, China (BGI-IRB E20007-T2).

## AUTHOR CONTRIBUTIONS

W.J., J.L., H.Y., H.C., and Z.O. conceived the study and designed the research. H.Y., J.L., X.X., W.J., X.J., B.F., X.C., Z.O., J.W., and H.Y. supervised the project. L.L., X.H., Y.W., X.L., W.L., N.Z., W.C., J.S., Y.W., J.Y., X.T., and H.Y. constructed the animal models, collected samples, and conducted IHC staining. P.R., J.L., T.Z., Y.Z., and Y.L. performed Stereo-seq experiments. A.C. and S.L. provided technical support for Stereo-seq experiments. G.H. conducted sequencing for Stereo-seq libraries. Z.O., P.R., T.Z., F.H., J.C., Q.C., X.H., Q.T., H.C., and Q.Z. carried out bioinformatic analysis. Z.O., P.R., L.L., T.Z., and F.H. wrote the manuscript.

## ACKNOWLEDGMENTS

This study was supported by National Key Research and Development Program (2022YFD1800200), Hatch Project of State Key Laboratory for Animal Disease Control and Prevention (SKLADCP2023HP05), Gansu Provincial Postdoctoral Special Project (23JRRA558), The Major Science and Technology Project of Gansu Province (23ZDNA007, 22ZD6NA001), Innovation Program of Chinese Academy of Agricultural Sciences (CAAS-ASTIP-2021-LVRI) and Natural Science Foundation of Qinghai Province (grant number 2021-ZJ-933). We thank China National GeneBank for providing sequencing services for this project. We would also like to express our appreciation to the computing platform STOmics Cloud (https://cloud.stomics.tech) for their support in enabling workflow automation and expediting the analysis of Stereo-seq data. Furthermore, we would like to acknowledge the efforts of Qingqing Yang, Fuling Xu, Guodong Dai, Wenying Lu, and Linsheng Zhang for their diligent work in collecting the samples. Additionally, we would like to thank Xinrui Cui, Qinglin Wang, and Yueying Huang for their assistance in the bioinformatics analysis.

## SUPPLEMENTAL INFORMATION

**Figure S1.**
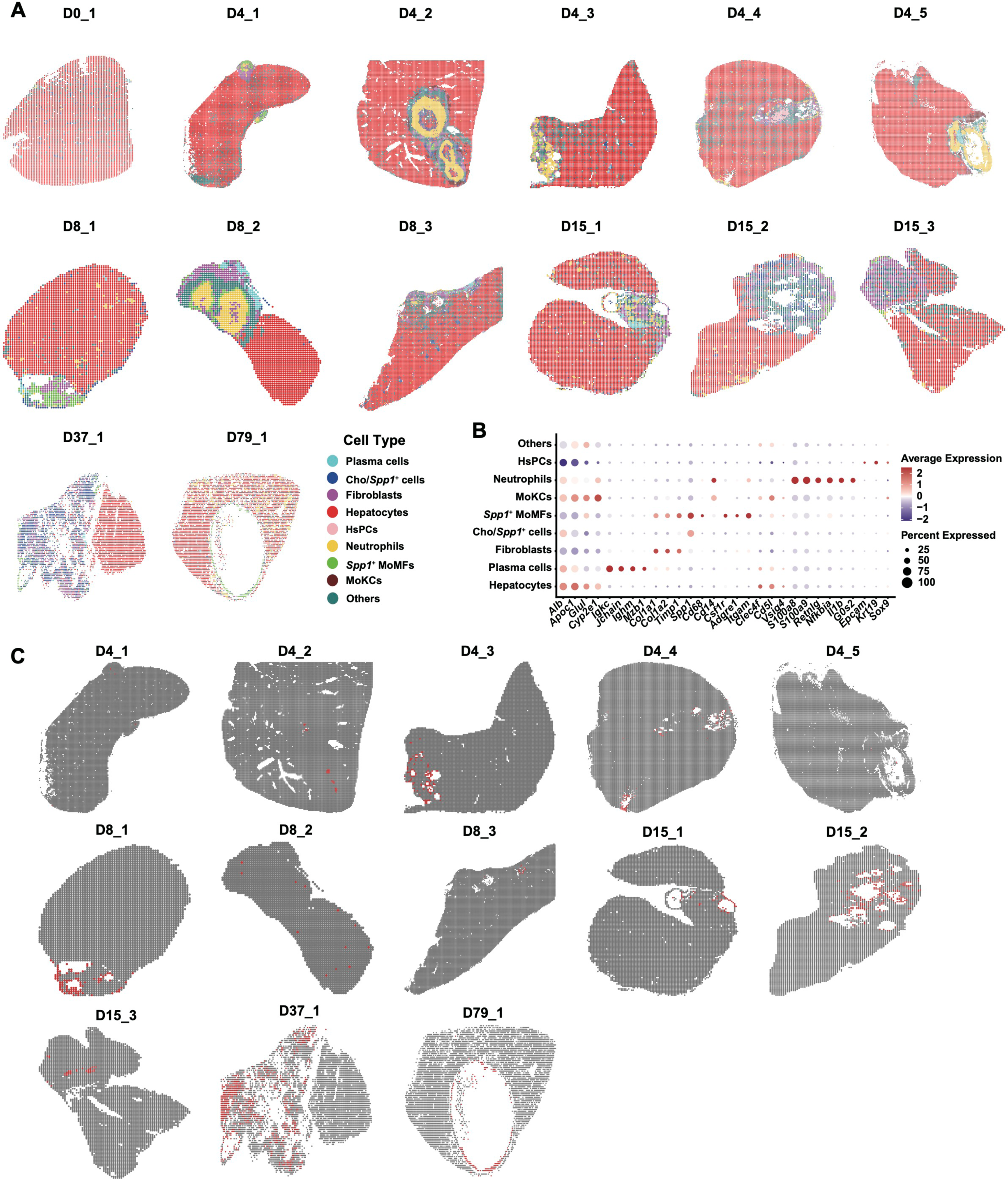
The spatial atlas of mouse livers with or without *Echinococcus multilocularis* infection; related to Figure 1. (A) Annotation results of 14 Stereo-seq chips. (B) Expression of the cell type markers in cell clusters identified based on Stereo-seq data. (C) Detection of *E. multilocularis* genes in Stereo-seq chips. No parasitic genes were detected in the control liver sample.

**Figure S2.**
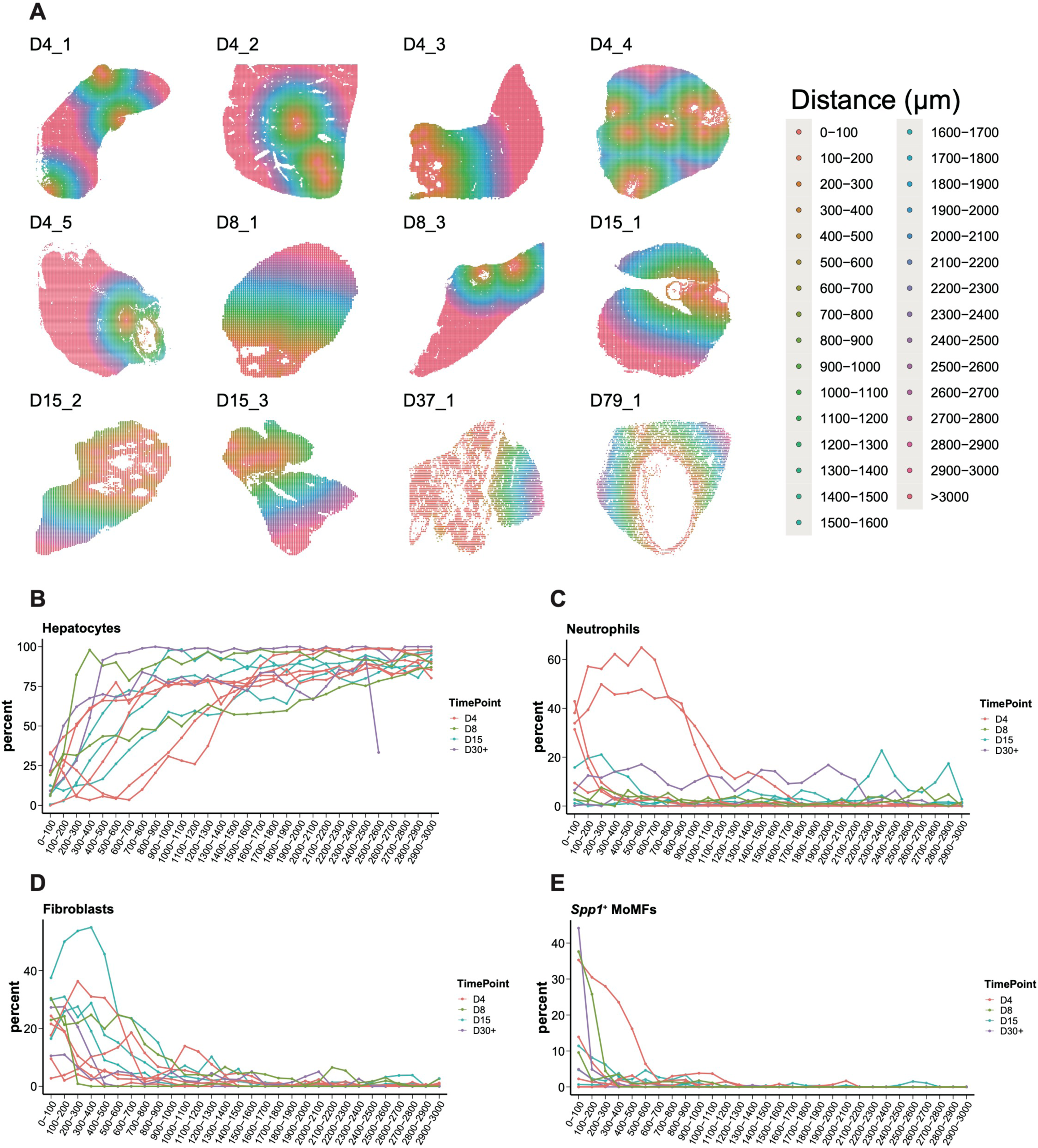
Spatial distribution of different cell types in mouse liver infected with *Echinococcus multilocularis*; related to Figure 1. (A) Circular expansion of the tissue layers based on a unit of 200 μm, starting from the AE lesion center. (B) Distribution of hepatocytes from the center to the distal region of the AE lesion in samples of different timepoints. (C) Distribution of neutrophils from the center to the distal region of the AE lesion in samples of different timepoints. (D) Distribution of fibroblasts from the center to the distal region of the AE lesion in samples of different timepoints. (E) Distribution of *Spp1^+^* MoMFs from the center to the distal region of the AE lesions in samples of different timepoints.

**Figure S3.**
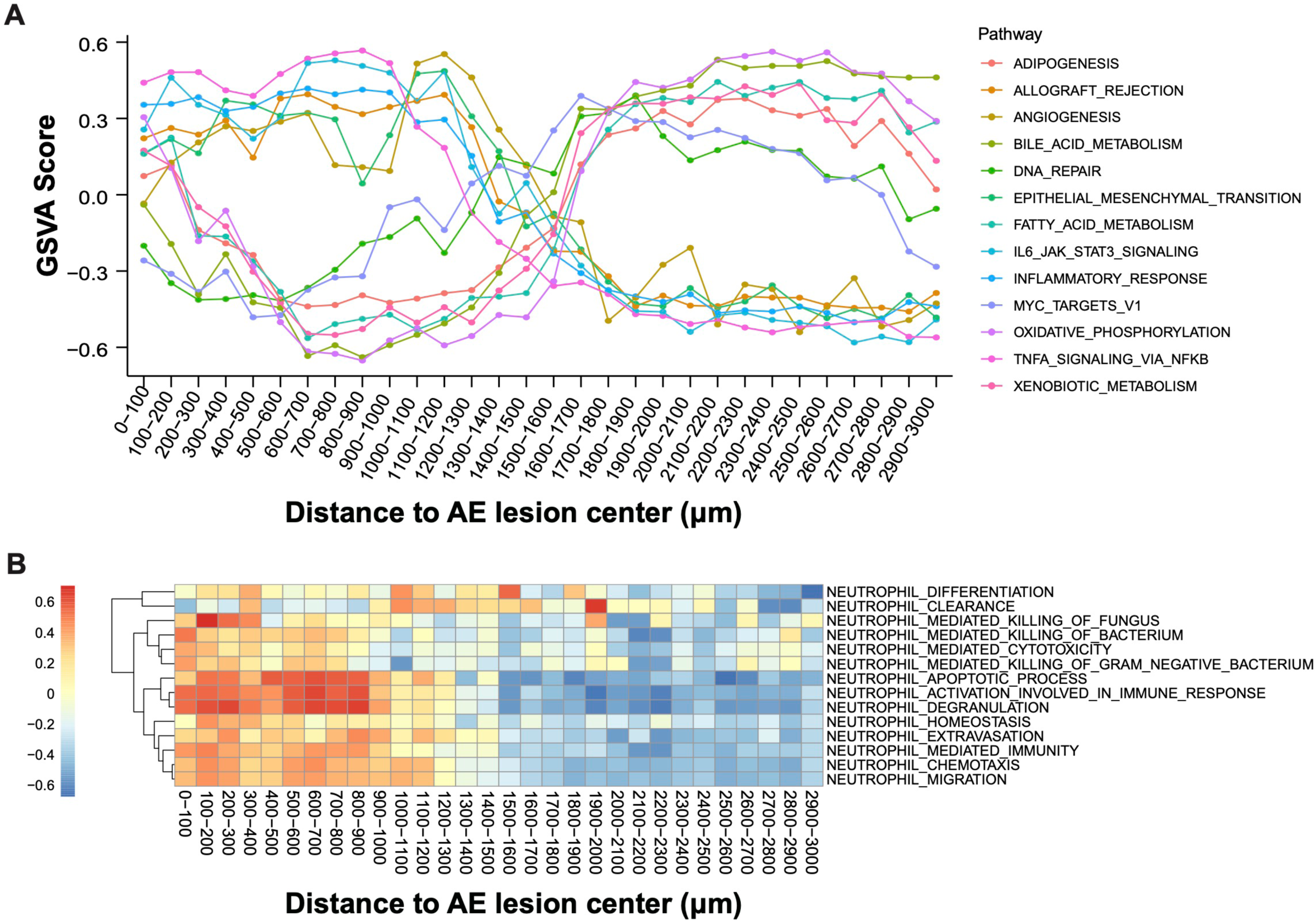
Spatial dynamics of biological pathways in AE lesion; related to Figure 1. (A) GSVA scores of multiple biological pathways from the center to the distal region of the AE lesion of Sample D4_1 (4 days post infection, Sample No.1). (B) GSVA scores of neutrophil-associated pathways from the center to the distal region of the AE lesion in Sample D4_1.

**Figure S4.**
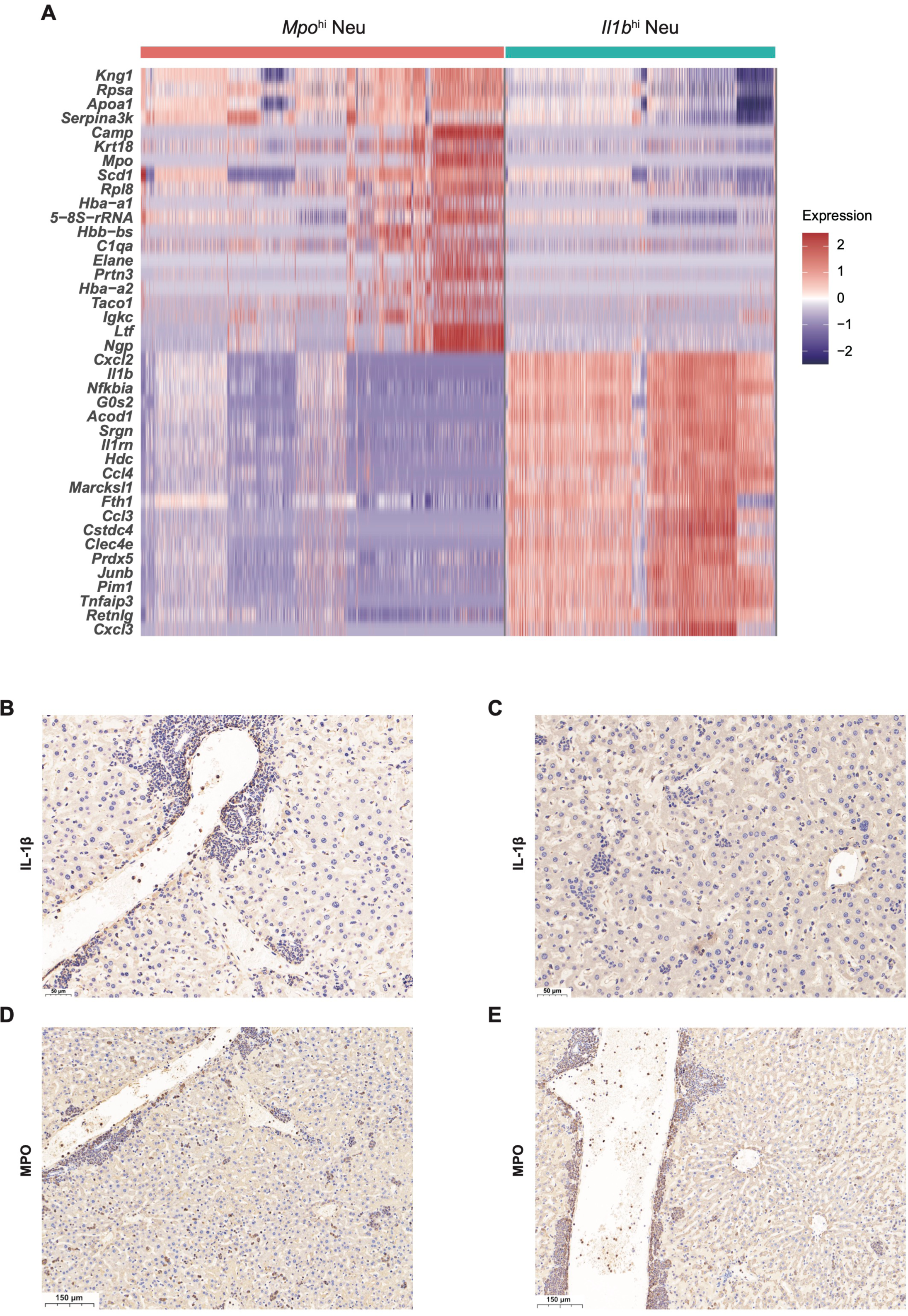
Identification and immunohistochemical staining of neutrophils in mouse livers infected with *E. multilocularis* (60 dpi); related to Figure 3. (A) Heatmap of differentially expressed genes in *Mpo^hi^* Neu and *Il1b*^hi^ Neu based on the Stereo-seq data. (B) Neutrophils adjacent to the AE lesion that were positive of IHC staining for IL-1β. (C) Neutrophils in the distal region of the AE lesion were negative of IHC staining for IL-1β. (D) Neutrophils adjacent to the AE lesion that were positive of IHC staining for MPO (myeloperoxidase). (E) Neutrophils in the distal region of the AE lesion were positive of IHC staining for MPO.

**Figure S5.**
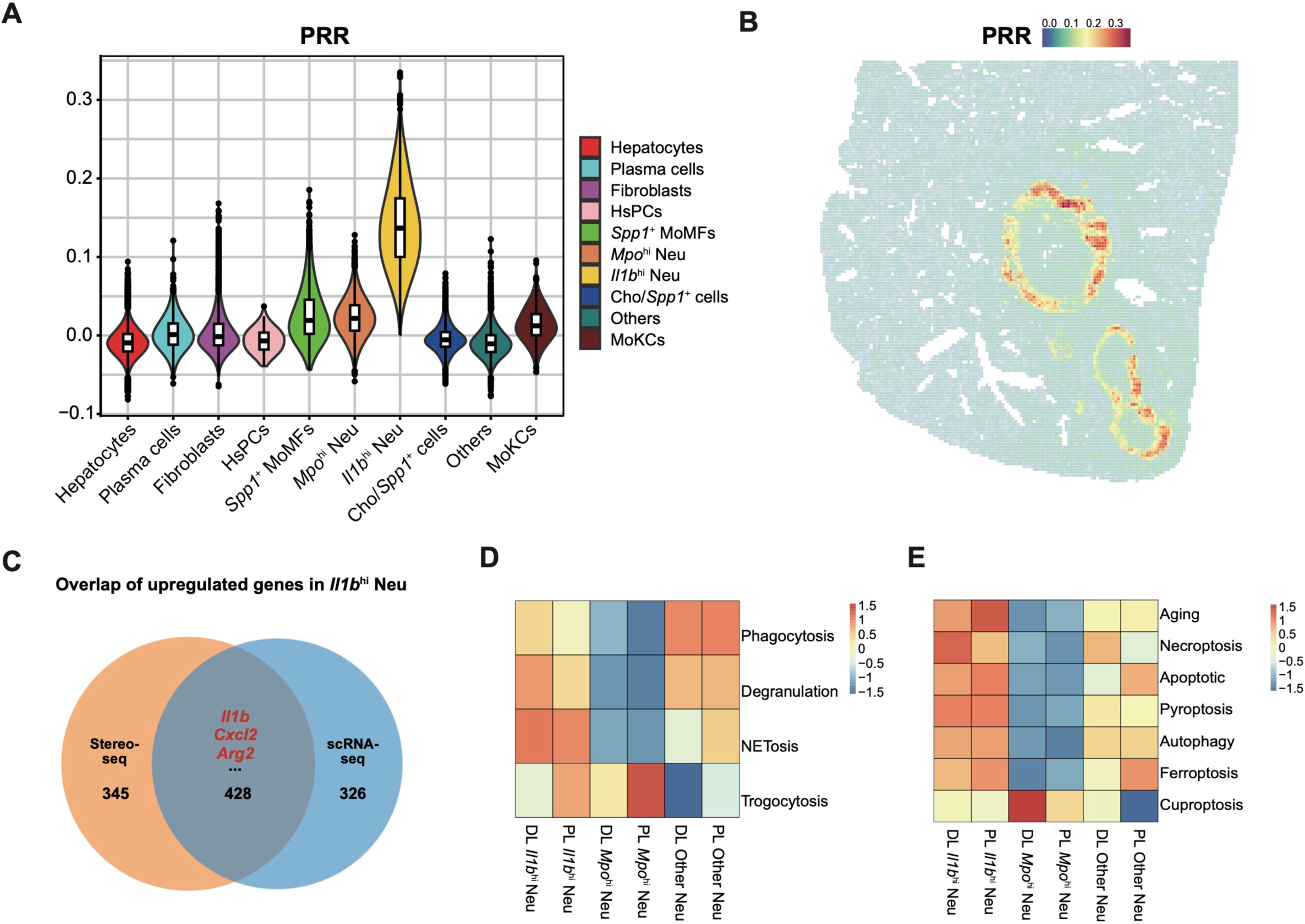
Transcriptional and functional heterogeneity of neutrophil subpopulations; related to Figure 3. (A) Comparison on the pattern-recognition receptor (PRR) gene set scores between all the cell types identified in 14 Stereo-seq chips. (B) Spatial pattern of PRR gene set signatures in Sample D4_2. (C) The overlap of upregulated genes in *Il1b*^hi^ Neu identified from Stereo-seq data and scRNA-seq data. (D) Heatmap showing the GSVA scores for pathogen-killing pathways of the neutrophil subgroups identified from scRNA-seq data. DL: distal-lesion; PL: peri-lesion. (E) Heatmap showing the GSVA scores for aging and cell death pathways of the neutrophil subgroups identified from scRNA-seq data.

**Figure S6.**
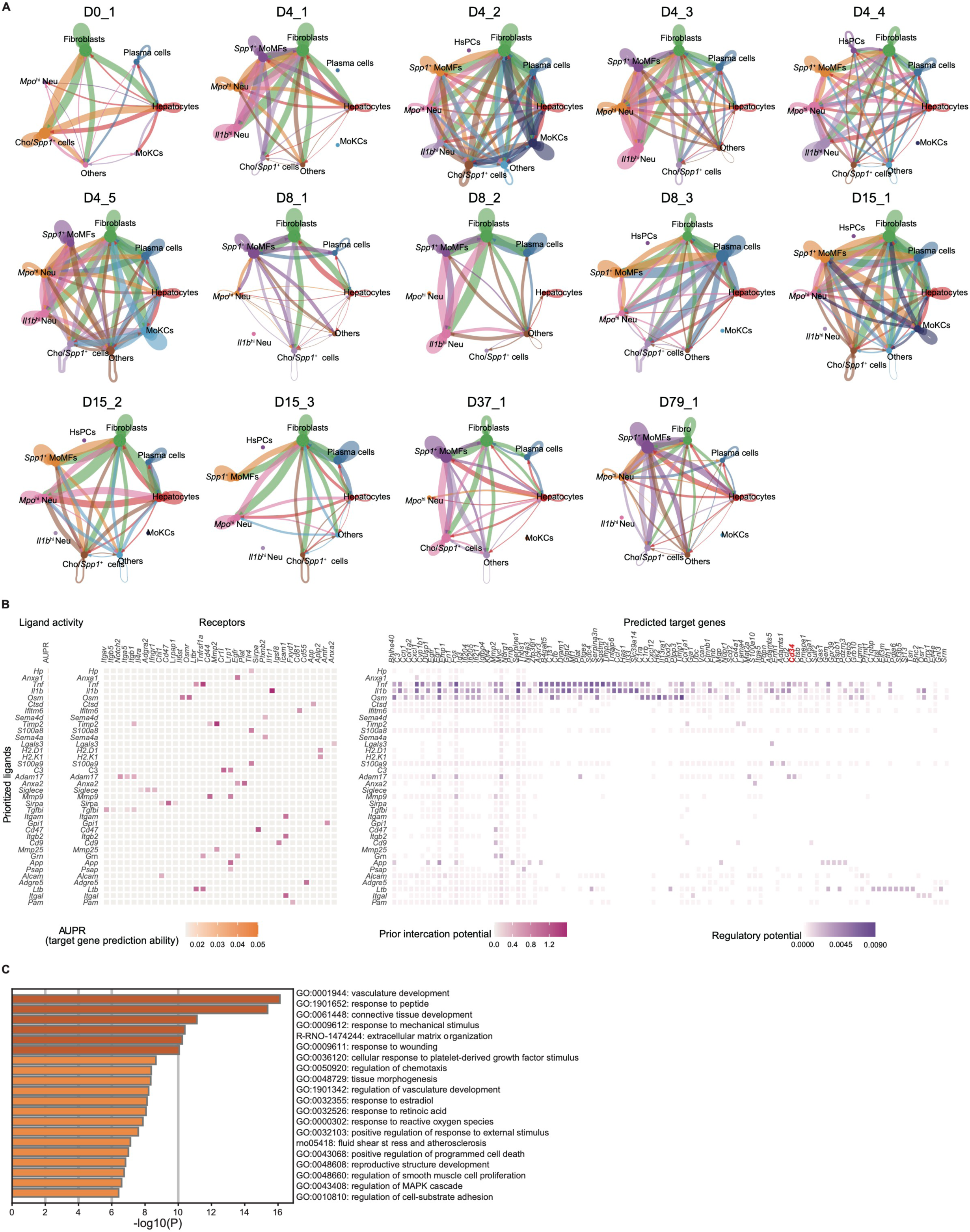
The cell-cell interactions between *Il1b*^hi^ Neu and the other cell types; related to Figure 4. (A) The cell-cell interaction network of 14 Stereo-seq chips. The line width represents the number of ligand-receptor pairs, and the arrow points to the receiver cell type. (B) Interactions between *Il1b*^hi^ Neu and fibroblasts identified by NicheNet, based on scRNA-seq data. Top-ranked ligands on *Il1b*^hi^ Neu are shown (left). Heatmap showing the ligand-receptor pairs between *Il1b*^hi^ Neu and fibroblasts arranged by ligand activity (middle). Heatmap showing the targets in fibroblasts that are potentially regulated by the ligands of *Il1b*^hi^ Neu (right). (C) GO enrichment of the target genes in fibroblasts that are potentially regulated by the ligands expressed by *Il1b*^hi^ Neu.

**Figure S7.**
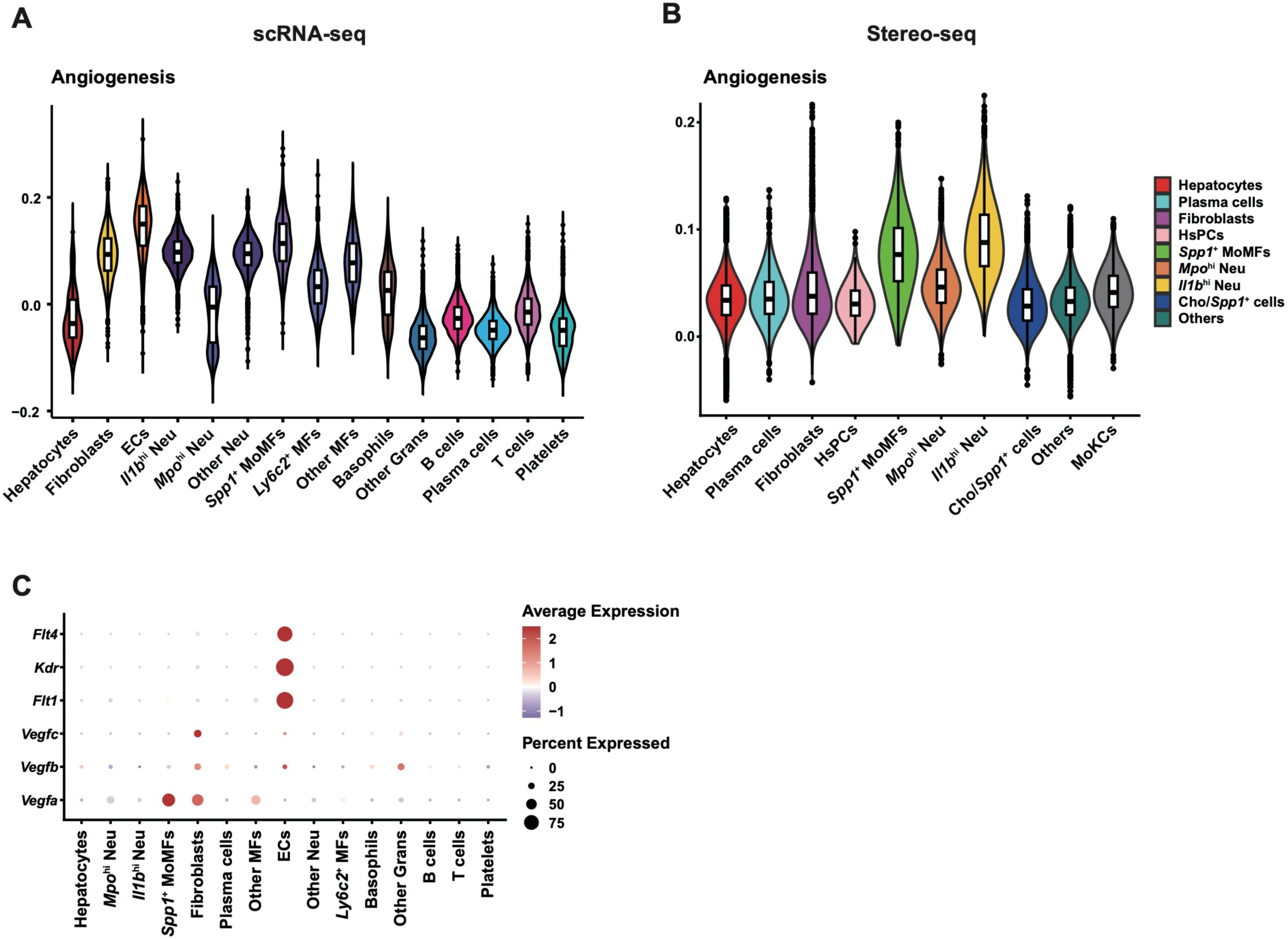
Angiogenesis potential of *Spp1^+^* MoMFs; related to Figure 5. (A) Angiogenesis gene set scores of all the cell types identified from the scRNA-seq data. (B) Angiogenesis gene set scores of all the cell types identified from the 14 Stereo-seq chips. (C) Dotplot showing the gene expression levels of *Vegfa*, *Vegfb*, *Vegfc*, and their receptors (*Flt1*, *Kdr*, and *Flt4*) based on the scRNA-seq data.

**Figure S8.**
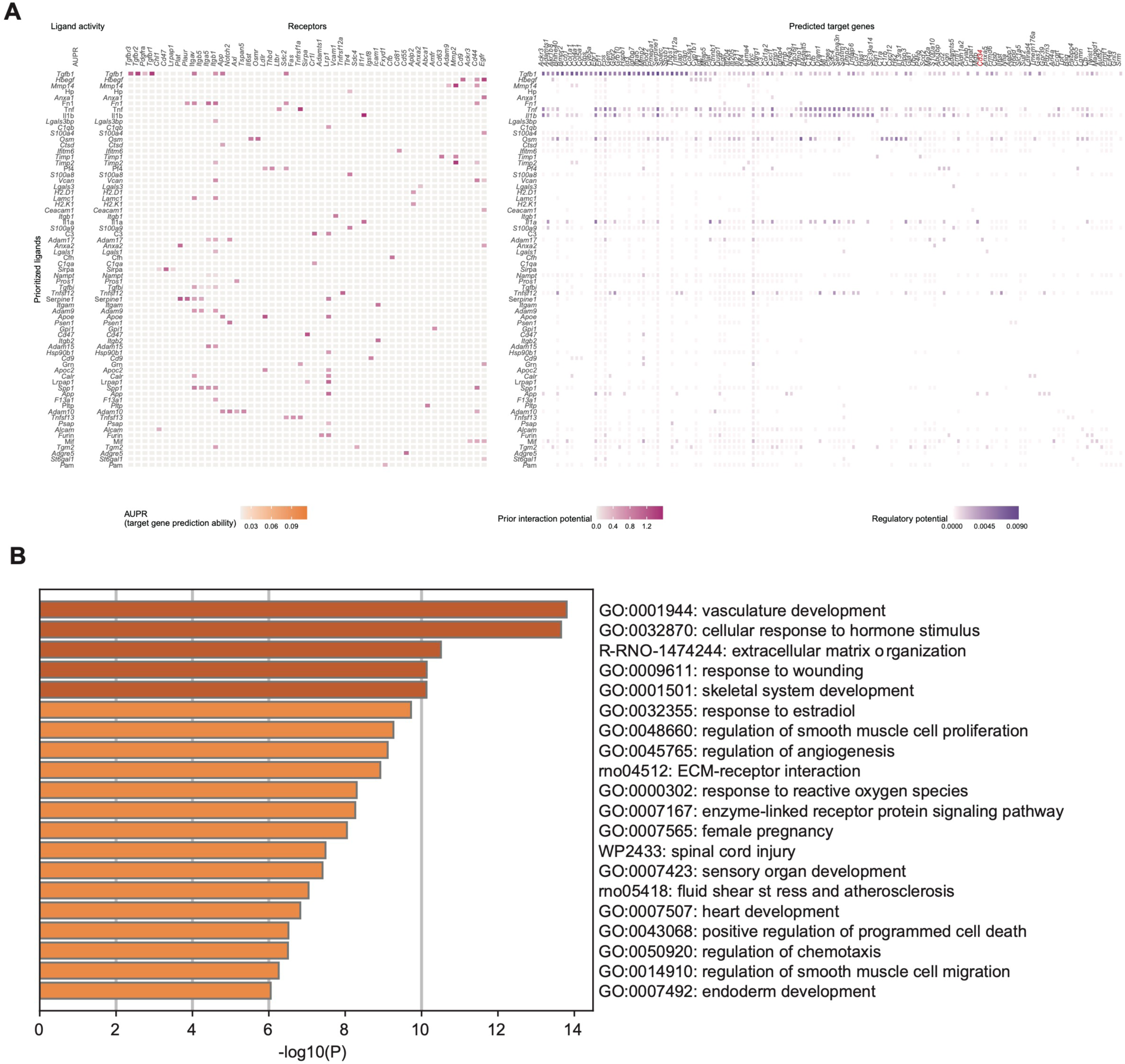
Ligand-receptor interactions between *Spp1^+^* MoMFs and fibroblasts identified from scRNA-seq data; related to Figure 6. (A) Top-ranked ligands on *Spp1^+^* MoMFs are shown (left). Heatmap showing the ligand-receptor pairs between *Spp1^+^* MoMFs and fibroblasts arranged by ligand activity (middle). Heatmap showing the targets in fibroblasts that are potentially regulated by the ligands from *Spp1^+^* MoMFs (right). (B) GO enrichment of the target genes in fibroblasts that are potentially regulated by the ligands from *Spp1^+^* MoMFs.

**Figure S9.**
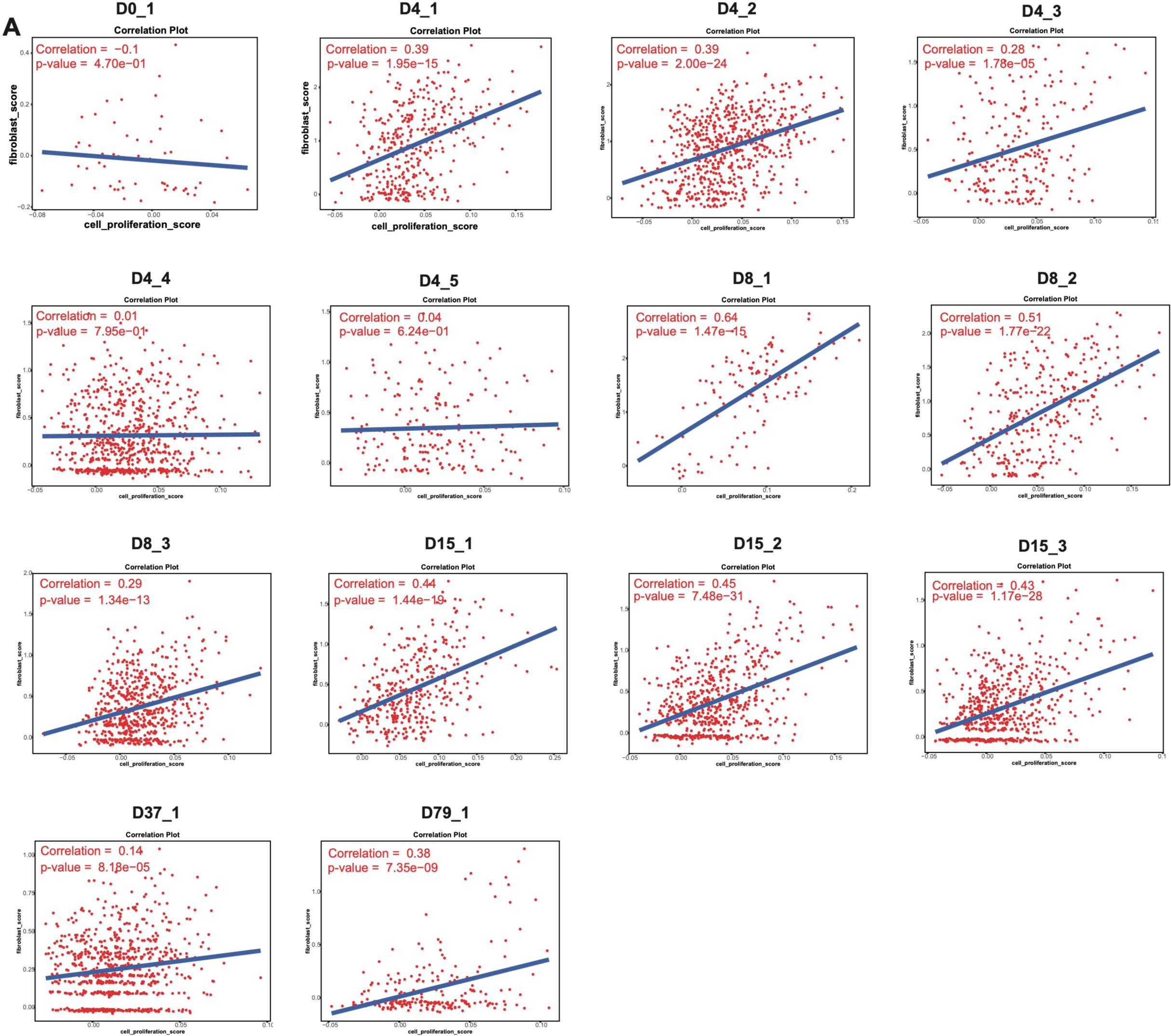
Assessment of the cell proliferation potential of fibroblasts; related to Figure 6. (A) Correlation analysis of fibroblast marker genes and cell proliferation features in 14 Stereo-seq chips.

**Table S1. Experimental details for the Stereo-seq samples.**

**Table S2. Annotation markers for the mouse scRNA-seq dataset**.

**Table S3. Gene sets used for GSVA and gene signature score analysis.**

